# LignAmb25: A Comprehensive AMBER Force Field Addressing Lignin’s Structural and Chemical Diversity

**DOI:** 10.64898/2026.01.03.697499

**Authors:** Marco Lapsien, Michele Bonus, Julian Greb, Holger Gohlke

## Abstract

LignAmb25 is a comprehensive force field for lignin molecular dynamics simulations implemented natively within the AMBER package. The force field includes parameters for all common monolignol units (*p*-coumaryl, coniferyl, caffeyl, and sinapyl alcohol) and their associated linkages (β-O4, β-5, β-β, β-1, 5-5, 5-O4, α-O4, BDO, and DBDO), along with less commonly encountered units such as tricin, spirodienones, and hydroxystilbenes. This enables simulations of both softwood and hardwood lignin structures with compositions that would be difficult to isolate experimentally. Force field parameters were initially derived from the GAFF2 force field and systematically optimized using quantum mechanical calculations at the ωB97X-D4/def2-TZVPP level of theory on conformer ensembles derived via the CREST/CENSO conformational sampling toolchain. Partial atomic charges were derived using the RESP methodology, consistent with AMBER conventions. Experimentally measured crystal structures of lignin simulated with LignAmb25 accurately retain their packing based on calculations of the RMSD and density error compared to the deposited crystal structure, thereby exceeding the performance of the lignin force field for CHARMM. Additionally, LignAmb25 is shown to reliably estimate the enthalpy of vaporization and the absolute hydration free energy of lignin-related compounds. The LignAmb25 force field is provided in two variants: LignAmb25^Solo^, a standalone version not meant for use with other biomolecular force fields that focuses on accurate modelling of lignin-solvent interactions, and LignAmb25^HF^, a version that is compatible with all other major biomolecular force fields in the AMBER molecular dynamics suite. This includes force fields of the GLYCAM (carbohydrates), ff19SB (proteins), and LIPID (lipids) families, as well as the DNA and RNA force fields routinely used in AMBER. The LignAmb25 force field will be distributed as of AMBER 26.

**Statement of significance:** Lignin, a complex aromatic heteropolymer comprising up to 40% of plant biomass, remains one of the most challenging biopolymers to characterize experimentally due to its structural heterogeneity, recalcitrance against depolymerization and selective chemical conversion, and lack of a defined primary sequence. Traditional wet-lab analytical methods face significant limitations, including lignin’s poor solubility, tendency to aggregate, and structural modifications during extraction and analysis. These experimental challenges make computational approaches essential for understanding the molecular basis of lignin’s physicochemical properties and for advancing lignocellulosic biomaterial applications. We present LignAmb25, a molecular mechanics force field for lignin implemented within AMBER, enabling researchers to investigate lignin structures and dynamics under conditions difficult to access experimentally. LignAmb25 integrates into the AMBER force field system and represents an alternative to the lignin force field for CHARMM. It improves upon the latter by including spirodienones and hydroxystilbenes as less commonly encountered monolignol units and the addition of several new linkages.

## Introduction

Valorization of biomass has gained increasing attention due to its abundance and the negative impact on climate of established procedures for obtaining raw materials (1, 2). Biomass is primarily composed of plant cell wall tissue, which itself is composed of the polysaccharides cellulose and hemicellulose, as well as lignin (3). The latter is a complex three-dimensional heteropolymer that provides structural integrity to the plant cell wall (4). Depending on the biomass source, lignin makes up to between 15-40% of the plant cell wall mass (5, 6), a noticeable fraction that is most often simply burned and, therefore, not properly valorized (7). Combating this waste of lignin is crucial, and depolymerization of lignin can yield sought-after functionalized aromatic compounds, such as vanillin, which is heavily used in the food industry (8), syringaldehyde, which is associated with the treatment of bacterial infections (9), or phenol, which can be used to build bio-based resins (10). Despite progress in valorization strategies, lignin’s full potential remains largely untapped. Deciphering its recalcitrance against depolymerization and selective chemical conversion is key to replacing petroleum-derived aromatics, such as benzene, toluene, and xylenes with renewable alternatives (11). Additionally, lignin plays a crucial role in cellulose-based textile production, where its presence negatively impacts yields (12) and spinning efficiency (13), both of which could likely be improved by a better understanding of lignin’s conformational dynamics and solvent interactions. Despite its potential, the molecular-level understanding of lignin’s behavior remains limited, hindering rational design of extraction processes, enzymatic degradation strategies, and lignin-based materials (14). The structural complexity of lignin presents unique analytical challenges that distinguish it from other biopolymers (15). Unlike proteins and nucleic acids, lignin lacks a defined primary sequence and exhibits substantial heterogeneity in both monomer composition and interunit linkage patterns due to the nature of its biosynthesis (16, 17). Lignin biosynthesis occurs via the coupling of oxidized monomer radicals and is therefore driven by stochastic variance (18).

Molecular dynamics (MD) simulations present a powerful approach to overcome the experimental limitations associated with lignin research and enable the investigation of lignin at atomic resolution. Computational studies have provided insights into lignin’s interactions with cellulose (19), conformational preferences of individual linkages (20), and passive transport across membranes (21). However, the accuracy of these simulations depends critically on the quality of the underlying force field parameters. While a lignin force field has been developed for CHARMM (22, 23), the AMBER (24) molecular dynamics package, widely used for biomolecular simulations, has lacked dedicated lignin parameters optimized using AMBER-consistent methodologies.

The development of accurate force field parameters for lignin presents unique challenges. The diversity of linkage types requires careful parameterization of numerous chemical environments that govern the three-dimensional structure (15, 25). To address this, we present LignAmb25, a comprehensive force field for lignin developed specifically for the AMBER molecular dynamics suite. Building upon the General Amber Force Field (GAFF2) (26) framework, we systematically optimized parameters using extensive quantum-mechanical (QM) calculations and conformational sampling via the CREST/CENSO (27, 28) conformational search pipeline. Our parameterization strategy focused on providing a modular force field that provides the, to date, most extensive set of linkages and monomers described in the literature. Furthermore, we prioritized integration with other existing biomolecular force fields implemented in AMBER, such as ff19SB (29), GLYCAM06 (30), LIPID21 (31), and the nucleic acid force fields (32, 33), routinely used to perform MD simulations.

The integration of LignAmb25 with existing AMBER force fields is particularly important for studying lignin-carbohydrate complexes (34), lignin-enzyme interactions (35), and the behavior of lignin in various solvent systems relevant to biorefinery processes (36). By providing these parameters to the community through the standard AMBER distribution, we aim to facilitate computational studies that can guide experimental efforts in lignin valorization and utilization.

## Materials and Methods

### Determination of force field scope

The choice of chemical species to include in the force field was based on experimentally observed monolignols and linkage types (Figure 1). This comprises the predominant coumaryl- (H), guaiacyl- (G), and syringyl- (S) lignin (15, 25), but also the less commonly found caffeyl-(C) lignin (37), and non-traditional monolignols such as the flavonoid tricin (38), as well as spirodienone (39) and hydroxystilbene (40) constructs. The varying chemical properties of the monolignols, coupled with the radical nature of the lignification process, lead to a variety of linkages found in lignin polymers. The β-O4, β-5, β-β, 5-O4, 5-5, and α-O4 linkages are among the more commonly encountered linkages (15, 25). However, less frequently encountered linkages, such as the benzodioxane (BDO) (41) and dibenzodioxocin (DBDO) (42) as well as α-cinnamate/-benzoate (43) and γ-cinnamate/benzoate (44, 45) linkages, were also included in the parametrization process. Additionally, linkages between lignin and carbohydrates were considered by including the β-O-C1, 4O-C1, α-O-C5, α-O-C2, and ferulate linkage types between monolignols and monosaccharides (43, 46).

**Figure 1:**
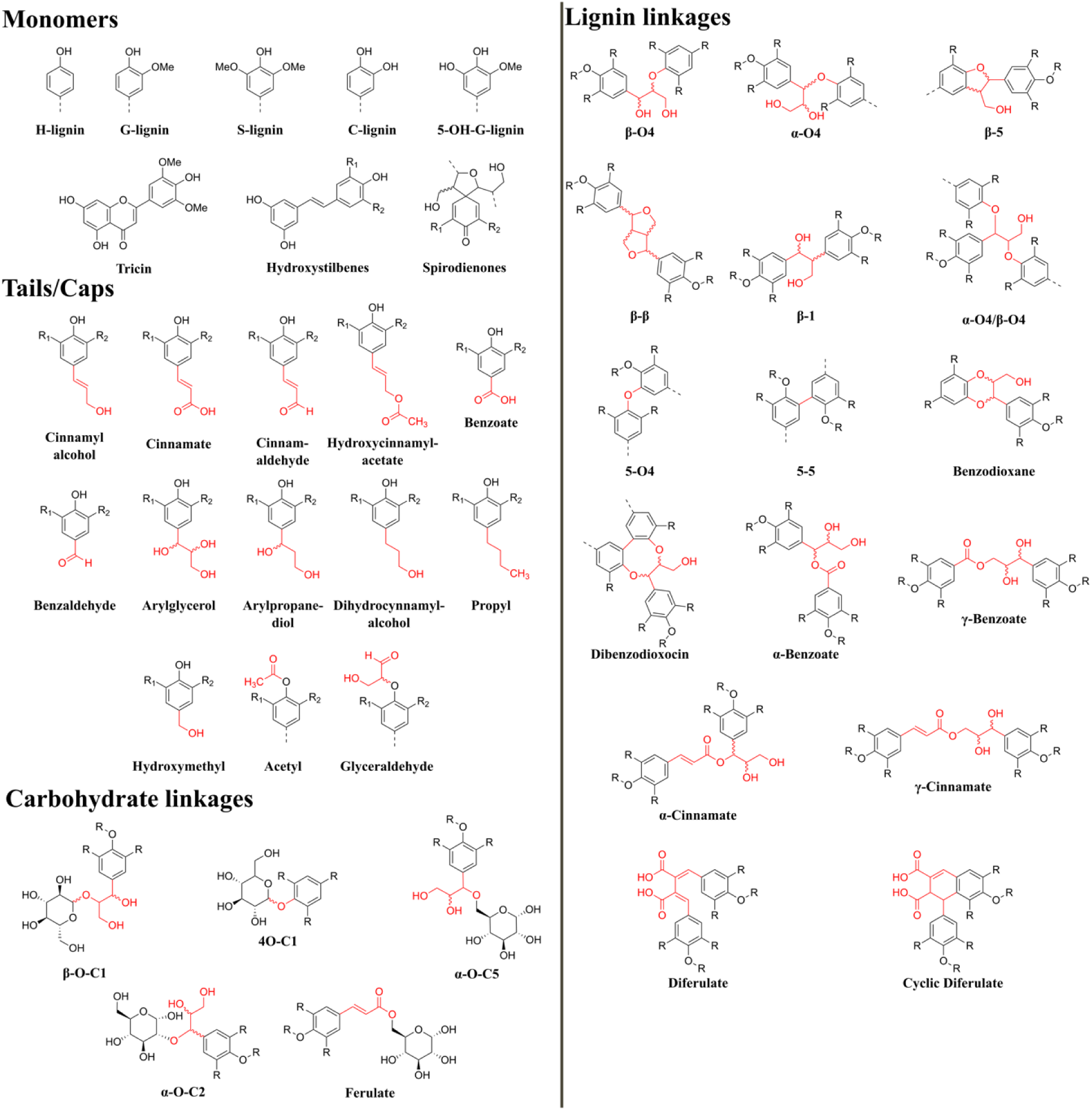
Overview of monomers, end groups, and linkages included in LignAmb25. Apart from abundant monolignols, the less common 5-OH-G-unit (37), tricin, hydroxystilbenes, and spirodienones were included. Each linkage, except for the spirodienone constructs, has dedicated fragments per stereoisomer (indicated by wavy bonds). Linkages between lignin and carbohydrates can exclusively be modeled with force fields from the GLYCAM family.

### Definition of force field fragments

Next, lignin dimers, trimers, and tetramers covering the pool of linkages included in the parametrization process were built, and a set of force field fragments was obtained by splitting the multimers at bonds connecting to the aromatic ring of a monomer (Figure 2). This approach was chosen as it reduces the number of fragments needed to model individual linkages, hence, keeping the total number of fragments reasonably low relative to lignin’s chemical complexity. This approach yielded distinct fragments for each linkage (with the exception of the 5-5 and 5-O4 linkage as a single bond separates the two aromatic rings) and separate monolignol fragments. In the case of H-, G-, S-, and C-lignin, they consisted only of the aromatic ring with a varying number and position of open valences depending on whether the ring is a branching point or part of a linear chain. Additionally, a variety of “tail” fragments was obtained that encapsulate the variations of monolignols at the C1 position. Although stereochemistry is often neglected in lignin research, a dedicated fragment of each stereoisomer per linkage was created, and in the case of the β-O-C1 and 4O-C1 lignin-monosaccharide linkages, variants specific to the α- and β-anomers were built.

**Figure 2:**
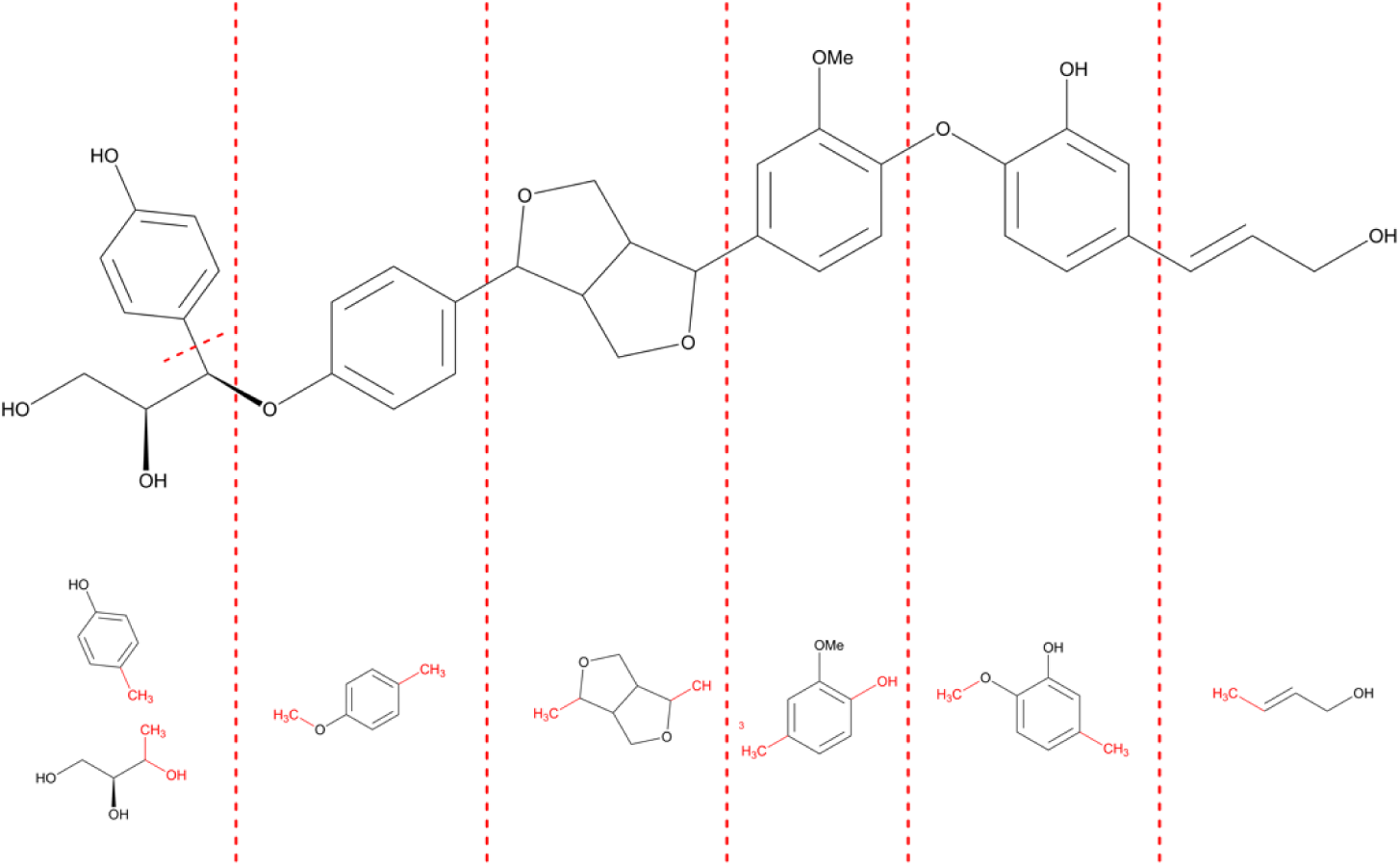
Capping strategy chosen for LignAmb25 development. The fragmentation of an example polymer into force field fragments is shown. Fragmentation points were chosen to maximize modularity of the force field while respecting established guidelines for RESP charge cap groups where possible (54). Most interunit linkages consist of ether bonds and carbon-carbon single bonds. Ether bonds were fragmented using a methyl group on the fragment carrying the oxygen and a hydroxyl group on the other fragment, and applying an intermolecular charge constraint of 0 between the two groups. Carbon-carbon single bonds were fragmented by attaching a methyl group to each fragment and applying an intramolecular charge constraint of 0 per capping group. As modularity of the force field definition was given a high priority, conjugated systems sometimes had to be fragmented, too (bottom right), which might impact the accuracy of charge derivation.

### Residue and atom type naming conventions

Residue names were chosen to avoid conflicts with existing biomolecular force fields in AMBER. This, unfortunately, limited the available three-character codes, resulting in non-trivial character combinations for most fragments. Aromatic fragments usually have the number of open valences as the first character, the type of monolignol as the second character, and a dedicated character that describes the position of the open valences, e.g., the code “1H1” corresponds to an H-unit with one open valence at the C1-position. Non-aromatic fragments mostly do not follow such a scheme and were instead assigned residue names not occupied by other force fields. Fragments with multiple stereoisomers share the same first two characters and differ in the third character to distinguish them. The letters “E” and “T” (in uppercase and lowercase, denoting one enantiomer each) represent the relative configurations *erythro*- and *threo*, where possible. Compounds with a single stereocenter will have “R” and “S” as the third letter in agreement with CIP nomenclature.

As atom types in the AMBER format only allow for two characters, the number of non-overlapping terms was even more sparse than for residue names (Table S1). The first character of an atom type corresponds to the element it describes (e.g., “O” for oxygen). Carbon atom types are an exception to this, as they are divided into aromatic (lowercase c) and non-aromatic (uppercase C) carbon atoms. The second character of each atom type was chosen to avoid overlaps with existing biomolecular force fields in AMBER.

### Conformational search

Representative ensembles of conformers for force field fragments and lignin multimers were derived using the Conformer-Rotamer Ensemble Sampling Tool/Conformer ENsemble Screening and Optimization (CREST/CENSO) conformational search pipeline. Initial models were generated using the RDKit Python package (version 2025.03.1) (47) and pre-optimized using the Geometry, Frequency, Noncovalent, second generation (GFN2) (48) method integrated into the extended Tight Binding (xTB) method (version 6.6.1) (49) in implicit water using the Generalized Born Surface Area (GBSA) solvation model (50). The resulting geometries were used as starting points for a conformational search using CREST (version 3.0.2) applying the Geometry, Frequency, Noncovalent Force Field (GFNFF) (51) method to avoid topology changes and the GBSA water model. The CREST conformer ensembles were subsequently filtered using CENSO (version 2.1.3) to obtain a representative, Boltzmann-weighted set of conformers. Details on the settings for each CENSO stage can be found in the Supporting Information (Table S2).

### RESP charge derivation

To obtain optimized partial atomic charges, a Restrained Electrostatic Potential (RESP) fit (52) was performed. Capping groups per fragment were chosen based on best practices recommended by the RESP ESP charge Derive (R.E.D.) server developers (53, 54). As a consequence of the approach chosen for defining force field fragments, they are primarily used to accommodate the splitting of ether and carbon-carbon single bonds (Figure 2). The former were split by attaching a hydroxyl capping group to one fragment and a methyl capping group to the other. To ensure compatibility with the GLYCAM force field family (30), the hydroxyl capping group was constrained to a charge of −0.1940, whereas the methyl capping group was constrained to the opposite charge of +0.1940. This ensures charge neutrality when lignin-carbohydrate linkages are built. The carbon-carbon single bonds were split by attaching a methyl capping group to each fragment and applying an intramolecular charge constraint of 0 to either capping group.

The Molecular Electrostatic Potential (MEP) per conformer of the CENSO-derived ensemble per force field fragment was calculated at the HF/6-31G* (55–57), MP2/cc-pVTZ (58, 59), B3LYP/6-31G* (60), B3LYP/cc-pVTZ, and ωB97X-D/cc-pVTZ (61) level of theory using Gaussian16 (62) with four shells built around the molecule in increments of 1.4, 1.6, 1.8, and 2.0 times the van der Waals radius according to the standard RESP procedure (52). The grid point density was increased to 17 points per Å^2^, and six different orientations per conformer were used to reduce the orientation dependence of the resulting RESP charge set (Figure S1) (63). The MEP of lignin multimers was calculated in the same manner, with only one orientation used per multimer.

The RESP fit was carried out using PyRESP (64) with the input files prepared using PyPE_RESP (65). Charge fitting was performed according to the standard two-stage procedure, where, in the second stage, only methyl groups were reoptimized. The additive RESP model with varying hyperbolic charge restraint weights (0.0003, 0.0005, 0.0007, 0.001) in the first stage and a fixed hyperbolic charge restraint of 0.001 in the second stage was applied. Conformers were fit together, with each conformer weighted according to the Boltzmann weight determined during the CENSO runs. Since six different orientations per conformer were used, the weight per orientation was equal to the conformer Boltzmann weight divided by 6. Mean RRMSE values obtained for RESP fits at different levels of theory and charge restraint weights are given in Table S3.

### Bond stretching parameter optimization

The lowest-energy conformer of each lignin multimer was used to perform potential energy surface (PES) scans using the r2SCAN-3c (66) composite method implemented in ORCA (version 6.0.0) (67). All bonds in each structure were scanned in steps of 0.02 Å in a range of ± 0.4 Å of the equilibrium bond length measured in the conformer geometry (41 steps total). Single-point energy calculations at the ωB97X-D4/def2-TZVPP (68–70) level of theory were performed on each scan geometry to obtain the final energy profile. Detailed information on convergence settings is provided in the Supporting Information (Table S4). PES energy profiles (*E*_*bond*_) were fit to the AMBER bond stretching potential energy term using the 17 points closest to the equilibrium (± 8 steps) (Eq. 1):

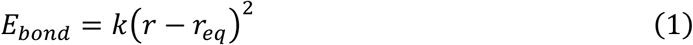

where *k* is the force constant, *r* is the scanned bond length, and *r*_*eq*_ is the equilibrium bond length. Bonds between identical atoms in different molecular contexts were grouped together based on atom types. Where necessary, these groups were further divided into subclusters using DBSCAN clustering based on force constant and equilibrium length variance, and a mean force constant and equilibrium length were derived for each (sub)cluster.

To assess improvement over GAFF2 parameters, single-point energy calculations on each PES scan geometry were performed using sander with the target bond term parameters nulled (1), changed to the mean cluster parameters (2), and changed to the GAFF2 parameters (3). The bond term contribution for the fitted and GAFF2 parameters was calculated by subtracting from (2) or (3) the single-point energies obtained with the bond stretching term nulled. The performance of the two parameter sets was assessed using the mean absolute error against the PES energy curve, with a decrease of at least 5% in the mean absolute error considered better and an increase of at least 5% considered worse.

### Angle bending parameter optimization

Parameters for angle bending were derived similarly to those for bond stretching, with PES scans performed in 2° increments over a range of ±40° from the equilibrium bond angle in the lowest-energy conformer geometry (41 steps total). Detailed information on convergence settings is provided in the Supporting Information (Table S4). The form of the objective function was according to the AMBER angle bending potential energy term (Eq. 2):

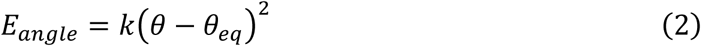

where *E*_*angle*_ is the potential energy, *θ* is the scanned bond angle, and *θ*_*eq*_ is the equilibrium angle. Improvement over GAFF2 parameters was assessed similarly to bond stretching parameters, including the previously fitted bond parameters in the parameter set.

### Torsion angle parameter optimization

Torsion angle parameters were derived by performing PES scans for each rotatable bond in each conformer of each lignin multimer in steps of 10° through a full 360°, starting from the equilibrium torsion angle measured in the conformer geometry (37 steps total). Detailed information on convergence settings is provided in the Supporting Information (Table S5). Torsional barriers (*V*), periodicities (*n*), and phase angles (*γ*) were derived using the Differential Evolution algorithm implemented in the evosax (version 0.2.0) (71) and JAX (72) Python packages with the objective function according to the AMBER torsion angle potential energy term (Eq. 3):

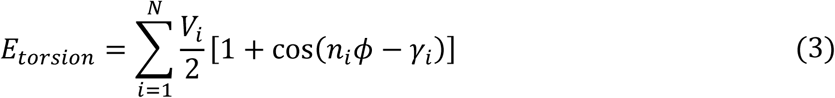

where *E*_*torsion*_ is the torsional energy, *N* is the maximum number of terms in the cosine series, and *ϕ* the scanned torsion angle. A maximum of *N* = 4 terms per dihedral pattern targeting the rotated bond was allowed. Additionally, dihedral patterns were weighted according to the number of potentials that arise per rotatable bond, and the angle values of all torsion patterns upon rotation of the bond were considered in the fit. Phase angles were constrained to multiples of 30° (i.e., 0°, 30°, 60°, 90°, 180°, or 270°). Optimization employed a hierarchical search strategy with an initial screening (15 generations, population size 25) followed by a full optimization (80 generations, population size 40) of the top 10% of parameter combinations. Early stopping was applied when no improvement was observed for 20 generations.

A mean parameter set per molecule was obtained by selecting the fitted parameter set yielding the lowest Boltzmann-weighted RMSE against the PES energy profiles of all conformers of the respective lignin multimer. The conformer-averaged parameter sets were then grouped by the atom types within the rotatable bonds. Subclusters were identified using hierarchical clustering with Ward Linkage on pairwise RMSE distances between Boltzmann-weighted mean energy profiles, with an RMSE threshold of 3.0 kcal mol^-1^ and a minimum cluster size of five members; smaller clusters were merged with their nearest neighbor cluster. The best parameter set within each subcluster was identified by selecting the fitted parameter set yielding the lowest Boltzmann-weighted RMSE against the PES energy profiles of all conformers of all molecules in the cluster.

Improvement over GAFF2 parameters was assessed similarly to bond stretching and angle bending parameters, including the previously fitted bond and angle parameters in the parameter set.

### Unbiased molecular dynamics simulations

All MD simulations described in this work were performed using the baseline version (gas-phase simulations), MPI-accelerated (minimization and equilibration), and GPU-accelerated version (production simulations) of the pmemd molecular dynamics engine as part of AMBER24 (24, 73). Simulations were done using either the newly developed LignAmb25 force field or the General Amber Force Field (GAFF2). Partial charges associated with the GAFF2 force field were derived using the RESP protocol described above with MEPs calculated at the HF/6-31G* level of theory.

For additional details on relaxation, thermalization, and production simulations per system, see the respective section below. Trajectories were analyzed using cpptraj (74).

### Modeling and simulation of crystal systems

18 crystal structures of lignin-related compounds and lignin dimers were taken from the Cambridge Structural Database (75). The structures were previously simulated using the lignin force field for CHARMM (LF-CHARMM) and therefore allow direct comparison between LF-CHARMM and LignAmb25 (23). Unit cells of the crystals were taken as provided from the simulations using LF-CHARMM. The PropPDB and ChBox tools from AmberTools25 were used to extend unit cells to 50.0 Å in each dimension and to ensure that experimental unit cell dimensions are retained (24).

Three minimization steps with gradually decreasing harmonic restraints from 25.0 to 15.0 to 10.0 kcal mol^−1^ Å^−2^ on heavy atoms were performed using the steepest descent algorithm for 2500 steps and the conjugate gradient algorithm for another 2500 steps (5000 steps total). The system was then thermalized over 200 ps from 80 K to the temperature at which the experimental data were collected under NVT conditions with harmonic restraints of 10 kcal mol^−1^ Å^−2^ applied to heavy atoms. Temperature scaling was controlled using Langevin dynamics with a collision frequency of 1.0 ps^−1^ (76). Six steps of 100 ps length each under NPT conditions were performed with gradually decreasing restraints on heavy atoms (10.0 to 0.01 kcal mol^−1^ Å^−2^) to adapt the system density to a pressure of 1 bar. Pressure was controlled using isotropic position scaling via the Berendsen barostat with a relaxation time of 1 ps (77). The total thermalization and density adaptation time was 800 ps. Production simulations were run for 20 ns over four independent replicas, with the 200 frames obtained from the last 10 ns of simulation time used for the calculation of mean crystal properties. The simulation time was chosen to be identical to the crystal simulations performed using LF-CHARMM to ensure comparability.

### Calculation of the enthalpy of vaporization

To estimate enthalpies of vaporization, gas-phase simulations were conducted by placing a single copy of the respective molecule in the simulation box with no periodic boundary conditions applied and a long-range interaction cutoff of 99.0 Å set. The liquid state was mimicked by placing 500 copies of the respective molecule in a 60×60×60 Å cube using packmol (78) with periodic boundary conditions and a real space interaction cutoff of 10.0 Å applied. Energy minimization was performed in three stages of 10,000 cycles each (8,000 steepest descent followed by 2,000 conjugate gradient), with harmonic restraints on non-hydrogen atoms progressively reduced from 25.0 to 15.0 to 10.0 kcal mol^−1^ Å^−2^. Thermalization and pressure adaptation consisted of an initial 100 ps NVT simulation with heating from 100 to 298 K, followed by six 100 ps NPT stages (700 ps total time) to adapt the system’s density to a pressure of 1 bar. Positional restraints were gradually decreased from 10.0 to 0.01 kcal mol^−1^ Å^−2^ over the six NPT stages. Temperature was maintained at 298 K using a Langevin thermostat with a collision frequency of 2.0 ps^−1^, and pressure was controlled using the Berendsen barostat with isotropic position scaling and a relaxation time of 1.0 ps for liquid-phase simulations. In both the gas-phase and liquid-phase simulations, all bonds involving hydrogen atoms were constrained using the SHAKE (79) algorithm, permitting a 2 fs integration timestep. Production simulations were carried out for 10 ns over four independent replicas, with Δ*H*_vap_ calculated via Eq. 4:

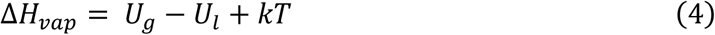

where *U_g_* and *U*_*l*_ represent the average molecular potential energies in the gas and liquid phase, respectively. *U_g_* and *U*_*l*_ were obtained by computing the average of the average molecular potential energy per replica.

### Calculation of hydration free energy

Absolute hydration free energies were calculated using a previously developed thermodynamic integration (TI) single-topology softcore annihilation approach (80). Solutes were solvated in TIP3P (81) water boxes (16 Å buffer), and decoupling was sampled in eleven *λ* windows determined using Gauss-Legendre quadrature nodes on [0, 1], where *λ* = 0 corresponds to full decoupling and *λ* = 1 to full coupling. Minimization and thermalization were performed at *λ* = 0.5 to provide geometries compatible with all windows. Production simulations of 5 ns per window (after 0.5 ns equilibration) were performed at 298 K using Langevin dynamics. Free energies were computed by numerically integrating ⟨∂*H*/∂*λ*⟩ using alchemlyb (82), with uncertainties estimated from four independent replicas. Experimental data were taken from the FreeSolv Database (83).

## Results & Discussion

### Partial charge assessment

Since the LignAmb25 force field is founded upon GAFF2, all parameters except partial atomic charges could be used as a starting point. RESP fits on the capped force field fragments were performed using MEPs calculated at five different levels of theory (HF/6-31G*, B3LYP/6-31G*, B3LYP/cc-pVTZ, ωB97X-D/cc-pVTZ, and MP2/cc-pVTZ), applying multiple charge restraint weights, to obtain partial atomic charges (Figure 3). The quality of MEP reproduction, calculated at the same level of theory, was tested with regard to the RESP fits per fragment (Figure 3A), as well as for molecules assembled from RESP charge sets for fragments (Figure 3B). The comparison emphasized the robustness of RESP fits, as all levels of theory showed excellent results for most fragments with a mean RRMSE value of ca. 0.09 (Table S3). The similarity of fit results is further reflected in the distribution of outliers, which were nearly identical for all levels of theory tested. These outliers were identified as fragments with vicinal hydroxyl groups, which appear to create a MEP landscape that is difficult to capture using point charges. This hypothesis is further supported by the excellent correlation between fits with and without cap constraints, which showed *R*^2^ values ≥ 0.99 for all levels of theory (Figure S2), suggesting that the additional constraints do not cause the outlier behavior.

**Figure 3:**
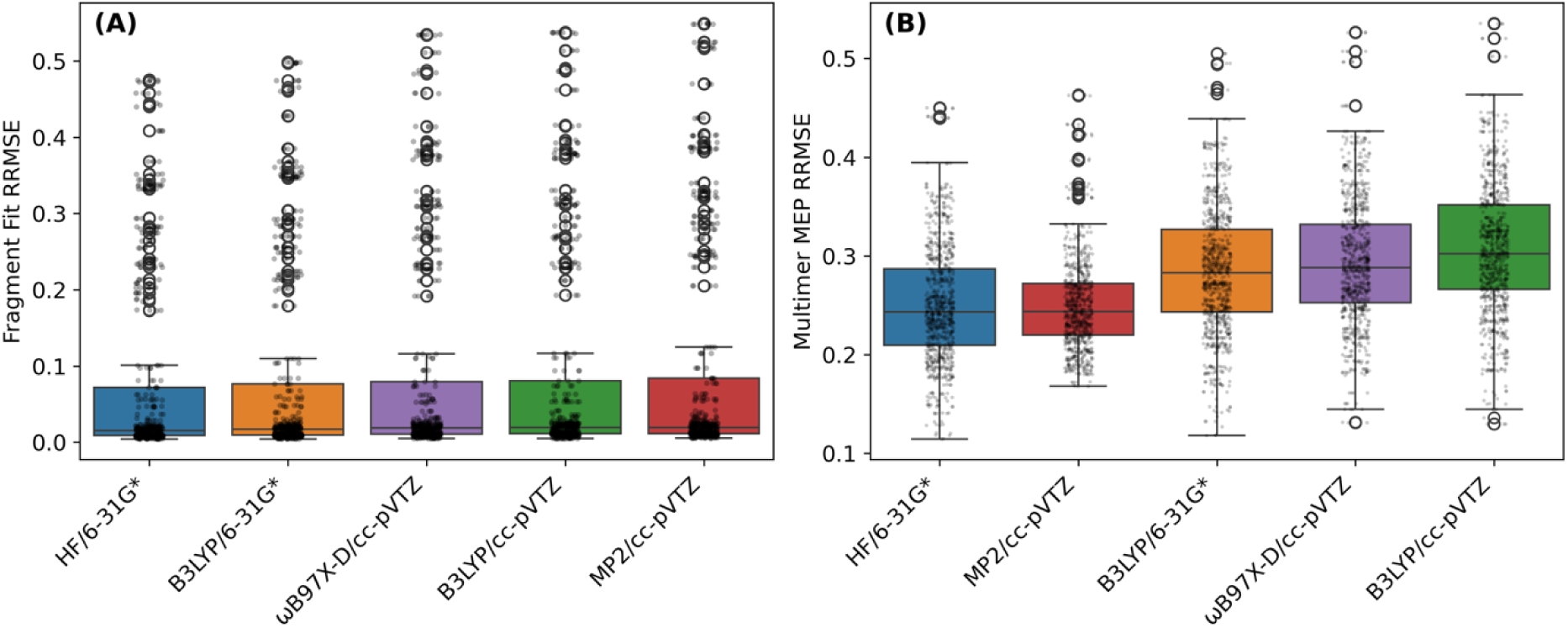
Quality of RESP charge fitting across theory levels. (A) Distribution of fragment-level RRMSE of the comparison of MEP reproduced from RESP charge fits with quantum-mechanically calculated MEP at the respective level of theory. Each box represents the distribution across all fragments (*n* = 134) at the indicated theory level, with individual data points overlaid. (B) Distribution of MEP RRMSE for lignin multimers assembled from force field fragments against quantum-mechanically calculated MEP of the multimer at the respective level of theory. Each box represents the distribution across the three most populated (Boltzmann-weighted) conformers per multimer (*n* = 200). Box plots show median (center line), interquartile range (box), and 1.5× IQR (whiskers). Theory levels are ordered by mean value (best to worst, left to right). Combined data for charge restraint weights of 0.0003, 0.0005, 0.0007, and 0.001 are shown. Adjustment of charge restraint weights did not alter RRMSE distribution markedly (Figure S4).

Analysis of charge transferability to assembled molecules revealed that the DFT methods overall performed worse than the (post-)HF methods, with the HF/6-31G* and MP2/cc-pVTZ levels of theory yielding the best results, and B3LYP/cc-pVTZ yielding the worst. The MEP of conjugated systems, such as hydroxystilbenes and systems containing cinnamate and benzoate linkages was consistently found to be poorly reproduced for all investigated charge sets (Figure S3).

One can expect to obtain overall elevated RRMSE values when applying fragment charges in the context of a larger molecule. However, the elevation may have also been influenced by the definition of the force field fragment set, which was deliberately chosen to prioritize modularity over accuracy for single larger molecules by defining smaller, more general fragments. Consequently, this leads to disruption of conjugated systems, whose charge distributions cannot be properly captured when rebuilt from individual fragments. Furthermore, most capping groups on fragments were constrained to specific charges to ensure compatibility with GLYCAM, which removes further degrees of freedom.

### Optimization of bonded parameters

Initial evaluation of a subset of GAFF2 parameters on lignin multimers revealed substantial deviation from PES data, which made reparametrization necessary (Figure 4). RESP charge sets obtained with a charge restraint weight of 0.001 were used to optimize bonded parameters per level of theory, resulting in five candidate force fields (LignAmb25^HF/6-31G*^, LignAmb25^MP2/cc-pVTZ^, LignAmb25^B3LYP/6-31G*^,LignAmb25^B3LYP/cc-pVTZ^, and LignAmb25^ωB97X-D/cc-pVTZ^, named after the level of theory used for MEP calculation). The optimization protocol used PES scans of bonds, angles, and dihedrals for a set of model compounds covering various chemical environments present in lignin polymers (Figure S5). A single conformer per compound was used for optimizing bond stretching and angle bending parameters, whereas a Boltzmann-weighted set of conformers was used to parametrize torsions. Marginal improvement in terms of reproducing bond stretching energy profiles was achieved, as GAFF2 already covers this well. However, for angle bending profiles, the fitting protocol achieved improved reproduction of high-energy regions, lowering the RMSD across all profiles from 2.27 kcal mol^−1^ to 0.82 kcal mol^−1^. The high-energy regions are expected to be rarely sampled in simulations, but the improvement could be beneficial for dedicated sampling studies. The accuracy of torsion profile reproduction could be increased in low- and high-energy regions compared to GAFF2. The improvements also cover phase shifts and periodicity adjustments to match the QM-derived profiles (Figure 4). Note, however, that the GAFF2 terms were purposely defined to work on a variety of organic compounds, which required the use of “generic” terms that may be less suitable for a dedicated compound class.

**Figure 4:**
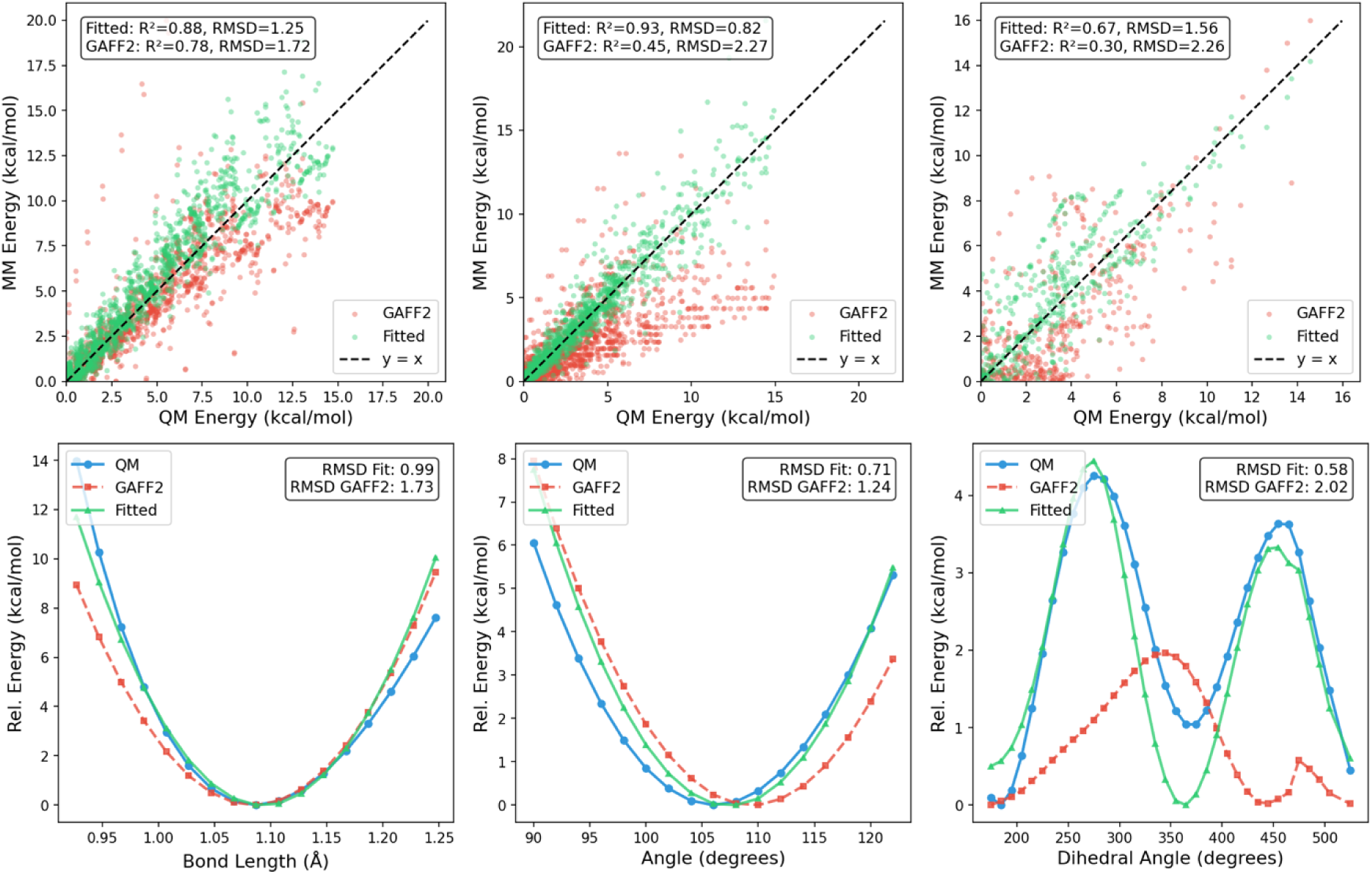
Comparison of fitted parameters of LignAmb25 with those from GAFF2 for bond, angle, and torsion terms at the B3LYP/cc-pVTZ level of theory. Top row: Parity plots comparing QM reference energies with MM energies for GAFF2 (red) and fitted parameters (green) for bond stretching, angle bending, and torsion parameters, respectively. The dashed line represents perfect agreement (*y* = *x*). Fitted parameters show improved correlation with QM for bonds (*R*^2^ = 0.88 vs. 0.78), angles (*R*^2^ = 0.93 vs. 0.45), and torsions (*R*^2^ = 0.67 vs. 0.30). Bottom row: Representative energy scans demonstrating improved agreement between fitted parameters and QM reference data: C-H bond stretch (left), H-O-C angle bend (middle), and C-C-O-H torsion rotation (right). In all cases, fitted parameters (green) reproduce the QM energy profile (blue) more closely than GAFF2 (red), with RMSD reductions of 43% (bond), 43% (angle), and 90% (torsion) for these examples.

### Thermodynamic validation

The optimized parameter sets per level of theory were used to compute enthalpies of vaporization (Δ*H*_vap_) and absolute hydration free energies (Δ*G*_hydr_) for a selection of lignin-related compounds (phenol, catechol, guaiacol, syringol, 4-hydroxybenzaldehyde, 4-propylguaiacol, 4-propylphenol) and small organic molecules that represent chemical environments found in lignin polymers (benzene, anisole, *m*-cresol, glycerol) to identify the best-performing parameter set based on MAE against experimental values (Figure 5).

**Figure 5:**
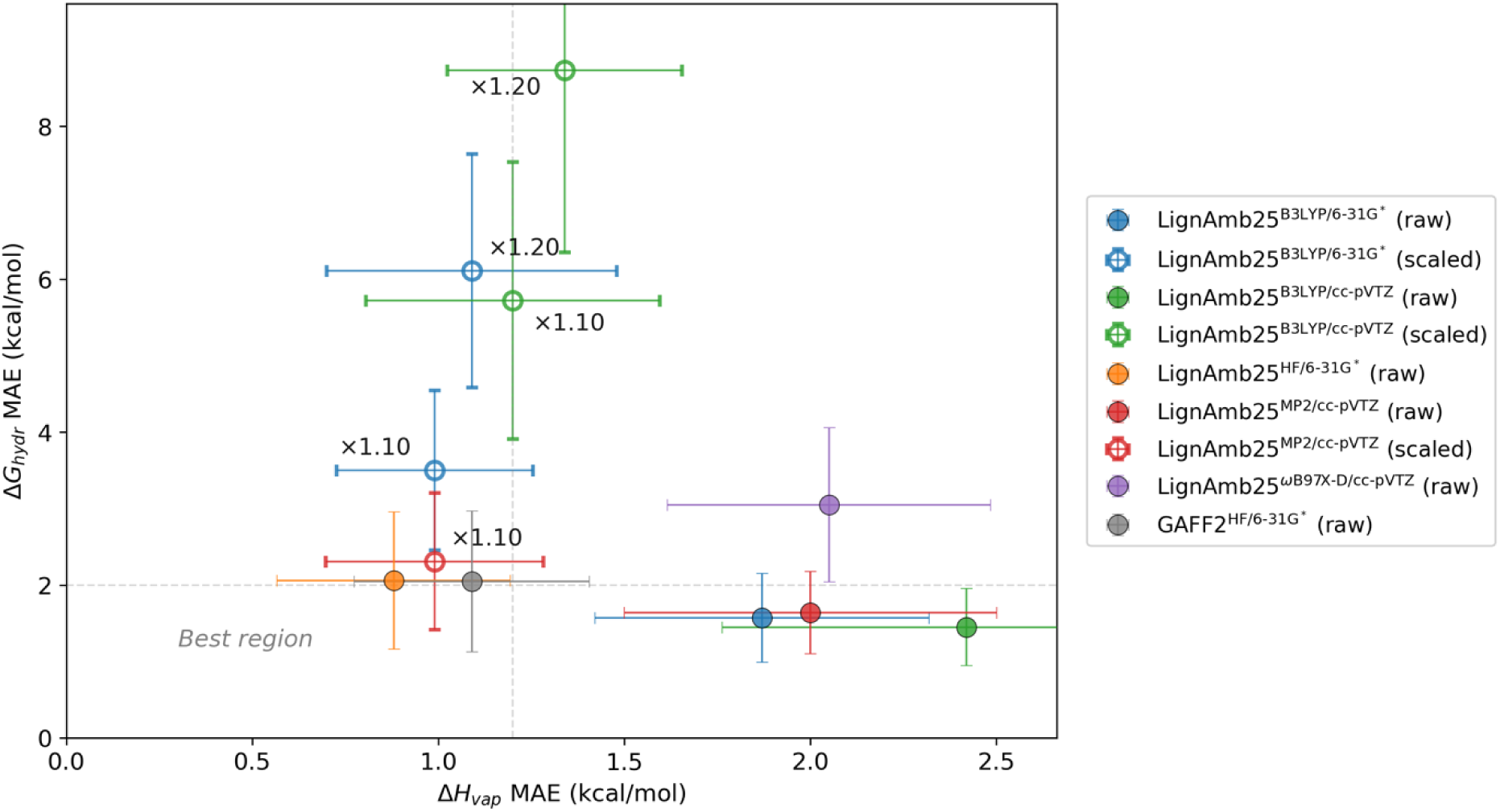
Trade-off between the prediction accuracy for heat of vaporization and hydration free energy. Each point represents a charge model, with Δ*H*_vap_ MAE (x-axis, *n* = 8 compounds) and Δ*G*_hydr_ MAE (y-axis, *n* = 11 compounds). Filled circles indicate raw (unscaled) charges; open circles indicate scaled charges (×1.10 to ×1.20). Colors denote the QM theory level used for MEP calculation. Dashed lines indicate approximate thresholds for acceptable accuracy chosen based on published force field benchmarks (85–87) (Δ*H*_vap_ MAE < 1.2 kcal mol^−1^, Δ*G*_hydr_ MAE < 2 kcal mol^−1^). Models in the lower-left region achieve good accuracy on both properties. Key models are labeled.

The GAFF2 bonded parameter set with HF/6-31G* derived charges (GAFF2^HF/6-31G*^) was used as a reference point and showed a good balance between MAE values of 1.10 ± 0.31 kcal mol^−1^ and 2.05 ± 0.92 kcal mol^−1^ for Δ*H*_vap_ and Δ*G*_hydr,_ respectively. LignAmb25^HF/6-31G*^ performed similarly, with MAE values 0.88 ± 0.31 kcal mol^−1^ and 2.06 ± 0.89 kcal mol^−1^. Interestingly, the LignAmb25^B3LYP/6-31G*^, LignAmb25^B3LYP/cc-pVTZ^, and LignAmb25^MP2/cc-pVTZ^ parameter sets behaved very similarly in that they predict well Δ*G*_hydr_ but give worse estimates for Δ*H*_vap_. LignAmb25^B3LYP/cc-pVTZ^ yielded the best prediction of Δ*G*_hydr_ (MAE = 1.45 ± 0.51 kcal mol^−1^), but at the same time the worst Δ*H*_vap_ prediction (MAE = 2.42 ± 0.66 kcal mol^−1^).

We experimented with rescaling B3LYP-derived charges by 1.10 and 1.20 to test whether the documented underestimation of dispersion effects in the B3LYP functional can be compensated for (84). Scaling of LignAmb25^B3LYP/cc-pVTZ^ charges by 1.10 considerably improved Δ*H*_vap_ prediction, lowering the MAE from 2.42 ± 0.66 kcal mol^−1^ to 1.20 ± 0.40 kcal mol^-1^ compared to the unscaled charges, but also degraded Δ*G*_hydr_ prediction (MAE = 5.72 ± 1.81 kcal mol^−1^ vs. 1.45 ± 0.51 kcal mol^−1^obtained with unscaled charges). A similar effect was observed when rescaling LignAmb25^MP2/cc-pVTZ^ charges, which decreased the Δ*H*_vap_ MAE to 0.97 ± 0.30 kcal mol^-1^ (surpassing GAFF2^HF/6-31G*^), but the Δ*G*_hydr_ MAE increased to 2.31 ± 1.03 kcal mol^-1^.

Ultimately, the unscaled LignAmb25^B3LYP/6-31G*^, LignAmb25^B3LYP/cc-pVTZ^, and LignAmb25^MP2/cc-pVTZ^ parameter sets were selected for further testing as they seemingly provide superior hydration free energy estimates than the balanced alternatives (GAFF2^HF/6-31G*^ and LignAmb25^HF/6-31G*^). We decided this because lignin is a ubiquitous plant-derived, highly heterogeneous structural biopolymer that is typically processed and utilized in dissolved or dispersed form, which will most likely be simulated more often in explicit solvent rather than as a solid or liquid. Therefore, an accurate representation of solvent-solute interactions is crucial.

Note, however, that mixing of DFT- or MP2- with HF-derived charges from other biomolecular force fields, such as the ff19SB protein force field (29), could lead to unbalanced interaction energies. We account for this by providing two versions of LignAmb25: LignAmb25^HF^ (shorthand for LignAmb25^HF/6-31G*^) that uses HF/6-31G*-derived RESP charges to ensure compatibility with other AMBER force fields, and LignAmb25^Solo^, which, based on crystal simulation results highlighted in the section below, was chosen to be LignAmb25^MP2/cc-pVTZ^ and is meant for standalone use if lignin-solvation questions are in the focus. Both versions use a custom set of atom types, as well as optimized bonded parameters obtained based on the respective charge set. Finally, note that the results in Figure 5 are specific to the use of the TIP3P (81) water model. The prediction of thermodynamic properties using other popular water models, such as OPC (88), needs to be assessed in future work.

### Crystal structure simulations

As a final comparison, 18 lignin-related crystal structures (Table S6) previously simulated using LF-CHARMM (23) were simulated using the GAFF2^HF/6-31G*^, LignAmb25^HF/6-31G*^, LignAmb25^B3LYP/6-31G*^, LignAmb25^B3LYP/cc-pVTZ^, and LignAmb25^MP2/cc-pVTZ^ force fields. RMSD of the unit cell against the experimental structures, as well as the density ratio of the experimental and simulated crystal structures, were computed and compared between the force fields (Figure 6).

**Figure 6:**
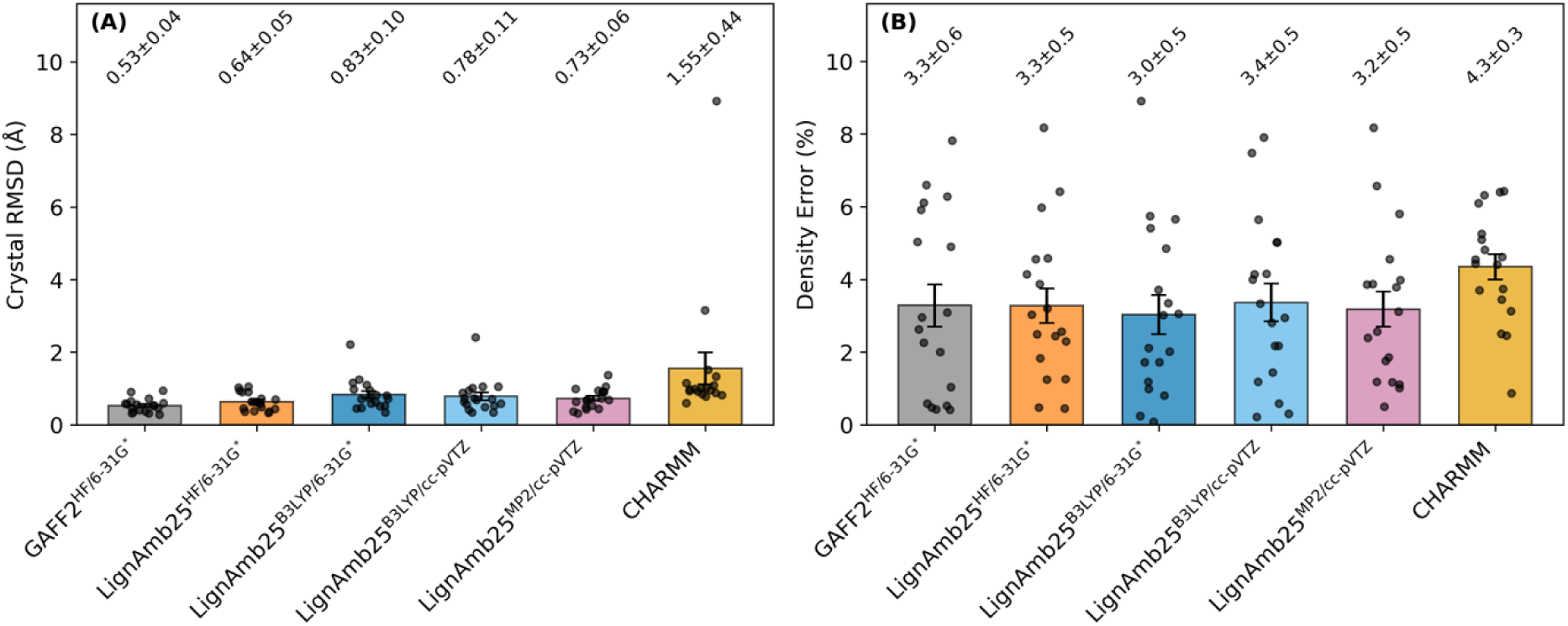
Crystal structure validation across force fields. (A) Mean absolute density error and (B) crystal heavy-atom RMSD for 18 lignin-related crystal structures (Table S6) validated in both this work and the CHARMM Lignin FF study (23). Data for GAFF2^HF/6-31G*^ and all LignAmb25 force field variants were obtained by taking the last 200 frames from the last 10 ns of a 20 ns-long MD simulation over 4 independent replicas. Bar heights represent mean values; error bars indicate standard error of the mean across structures. Density error calculated as |ρ_calc_/ρ_exp_ - 1| × 100%. Numerical data are available in the Supporting Information (Table S6).

GAFF2^HF/6-31G*^ performed best across all crystals with a mean density error of 3.3 ± 0.6% and a mean crystal RMSD of 0.53 ± 0.04 Å. All LignAmb25 force field variants showed comparable performance in terms of mean density errors of ca. 3.0%, but varying performance regarding mean crystal RMSD. LignAmb25^B3LYP/cc-pVTZ^ and LignAmb25^B3LYP/6-31G*^ showed RMSD values of 0.78 ± 0.11 Å and 0.83 ± 0.10 Å, respectively, and in either case, at least one crystal melted. The LignAmb25^HF/6-31G*^ set performed best out of the four LignAmb25 variants with a crystal RMSD of 0.64 ± 0.05 Å. LF-CHARMM also performed reasonably well, but worse than all LignAmb25 variants, with a mean density error of 4.3 ± 0.3% and a mean crystal RMSD of 1.55 ± 0.44 Å.

The results for GAFF2^HF/6-31G*^ underscore how well-behaved the GAFF2 parameter set is, but also that the fitted parameters used in the LignAmb25 force field variants yield results comparable to GAFF2^HF/6-31G*^ for most crystal systems and even improve upon LF-CHARMM in many cases. Since LignAmb25^MP2/cc-pVTZ^ was found to be most consistent in retaining crystal packing, it is chosen as the final LignAmb25 parameter set meant for standalone use. Note that the INELIW and INELIW01 systems contain two additional dioxane molecules. It can therefore not be ruled out that the dioxane parameters (in either case obtained from GAFF2) contribute unfavorably to system stability.

## Conclusion

We have developed LignAmb25, a molecular mechanics force field for lignin and lignin-related compounds with systematically derived partial charges that is available as both a standalone (LignAmb25^Solo)^ and AMBER-compatible (LignAmb25^HF^) version. LignAmb25 contains parameters for all common monolignol units and their associated linkages, along with less commonly encountered units such as tricin, spirodienones, and hydroxystilbenes. Through comprehensive evaluation of RESP charge fitting across five quantum mechanical theory levels, we identified B3LYP/cc-pVTZ as the best method for generating transferable charges that balance MEP reproduction with thermodynamic and structural accuracy. The chosen validation strategy revealed that MEP reproduction alone is an insufficient criterion for charge model selection. While all theory levels achieved good fragment-level fits (RRMSE < 0.10), thermodynamic validation exposed significant differences in predictive accuracy.

The LignAmb25^Solo^ parameter set yielded a MAE of 2.00 ± 0.50 kcal mol^-1^ for enthalpy of vaporization calculations, and hydration free energies with acceptable accuracy (1.64 ± 0.54 kcal mol^-1^) for the lignin-related compounds tested, while MAE values of 0.88 ± 0.31 kcal mol^-1^, and 2.06 ± 0.89 kcal mol^-1^ for enthalpy of vaporization, and hydration free energy calculations, respectively, were obtained using the LignAmb25^HF^ parameter set. Comparison with existing force fields demonstrates that LignAmb25 provides competitive performance. Crystal structure simulations on 18 structures common to both this work and the one describing the lignin force field for CHARMM showed comparable density reproduction across the force fields GAFF2^HF/6-31G*^ (3.3 ± 0.6%), LignAmb25^HF^ (3.3 ± 0.5%), LignAmb25^Solo^ (3.2 ± 0.5%), and LF-CHARMM (4.3 ± 0.3%). GAFF2^HF/6-31G*^ achieved the lowest crystal RMSD (0.53 ± 0.04 Å), followed by LignAmb25^HF^ (0.64 ± 0.05 Å), LignAmb25^Solo^ (0.73 ± 0.06 Å), and LF-CHARMM (1.55 ± 0.44 Å), though the LF-CHARMM results include one structure that exhibited crystal melting. Though GAFF2^HF/6-31G*^ showed remarkable performance, LignAmb25^HF^ delivered comparable results and was optimized to reproduce PES profiles more accurately in high-energy regions, which may be beneficial for extensive sampling studies.

Atom types and residue names in the LignAmb25 force field were carefully chosen to ensure compatibility with the major biomolecular force fields and GAFF2, allowing seamless integration into the AMBER MD Suite. We acknowledge that the large number of fragments in the force field and their unintuitive naming make structure preparation tedious. Future work will focus on developing a tool for automated preparation of lignin simulation systems, similar to what pdb4amber in AmberTools (24) provides for proteins. A prototype of this was used in this study to prepare most of the simulation systems. Finally, LignAmb25 is the most comprehensive lignin force field available to date, but it is by no means complete and could be extended in the future to include additional caps (e.g., methyl caps on hydroxyl groups) or tails (e.g., aminated tails) to capture the chemical diversity of lignin even more thoroughly and understand the role of these modifications better. The modular approach of LignAmb25 makes adding new residues straightforward. The LignAmb25 parameter and library files will be distributed with AMBER starting with AMBER26.

## Data and software availability

The LignAmb25 parameter and library files are available within the Amber suite of biomolecular simulation programs as of AMBER26. The Amber suite is available here: https://ambermd.org. Input files and analysis scripts for charge derivation and for computations of enthalpy of vaporization, absolute hydration free energy, and crystal simulations have been deposited at https://researchdata.hhu.de/ and are publicly available.

## Author contributions

HG designed the research. ML derived the force field parameters and performed validation. MB assisted in the modeling of electrostatic interactions and torsion parameters. JG assisted in choosing suitable methods for exploring conformational space. ML, MB, and HG wrote the manuscript. HG secured funding.

## Acknowledgements

This work was funded by a grant from the Ministry of Innovation, Science, and Research of North-Rhine Westfalia (NRW) within the framework of the NRW Strategieprojekt BioSC (No. 313/323-400-002 13) by the BOOST FUND 2.0 project “OptiCellu” and, in part, by the Deutsche Forschungsgemeinschaft (DFG, German Research Foundation) grant 270650915 (Research Training Group GRK 2158). We are grateful for the computational infrastructure and support provided by the “Zentrum für Informations- und Medientechnologie” (ZIM) at Heinrich Heine University Düsseldorf and the computing time provided by the John von Neumann Institute for Computing (NIC) to HG on the supercomputer JUWELS at Jülich Supercomputing Centre (JSC) (user IDs: VSK33, lignin). We acknowledge the developers of AMBER and the RDKit cheminformatics package for making their software publicly available.

## References

1. Höök M, Tang X. Depletion of fossil fuels and anthropogenic climate change—A review. Energy Policy. 2013;52:797–809.

2. Johnsson F, Kjärstad J, Rootzén J. The threat to climate change mitigation posed by the abundance of fossil fuels. Climate Policy. 2019;19(2):258–74.

3. Keegstra K. Plant Cell Walls. Plant Physiology. 2010;154(2):483–6.

4. Kang X, Kirui A, Dickwella Widanage MC, Mentink-Vigier F, Cosgrove DJ, Wang T. Lignin-polysaccharide interactions in plant secondary cell walls revealed by solid-state NMR. Nature Communications. 2019;10(1):347.

5. Olatunji KO, Ahmed NA, Ogunkunle O. Optimization of biogas yield from lignocellulosic materials with different pretreatment methods: a review. Biotechnology for Biofuels. 2021;14(1):159.

6. Saini JK, Saini R, Tewari L. Lignocellulosic agriculture wastes as biomass feedstocks for second-generation bioethanol production: concepts and recent developments. 3 Biotech. 2015;5(4):337–53.

7. Chio C, Sain M, Qin W. Lignin utilization: A review of lignin depolymerization from various aspects. Renewable and Sustainable Energy Reviews. 2019;107:232–49.

8. Fache M, Boutevin B, Caillol S. Vanillin Production from Lignin and Its Use as a Renewable Chemical. ACS Sustainable Chemistry & Engineering. 2016;4(1):35–46.

9. Ibrahim M, Sriprasanthi RB, Shamsudeen S, Adam F, Bhawani S. A concise review of the natural existence, synthesis, properties, and applications of syringaldehyde. Bioresources. 2012;7:2820–32.

10. Gong X, Meng Y, Lu J, Tao Y, Cheng Y, Wang H. A Review on Lignin-Based Phenolic Resin Adhesive. Macromolecular Chemistry and Physics. 2022;223(4):2100434.

11. Gong WH. BTX from Lignin. Industrial Arene Chemistry 2023. p. 1859–907.

12. Chen C, Duan C, Li J, Liu Y, Ma X, Zheng L, et al. Cellulose (dissolving pulp) manufacturing processes and properties: A mini-review. BioResources. 2016;11:5553–64.

13. Protz R, Lehmann A, Bohrisch J, Ganster J, Fink HP. Solubility and spinnability of cellulose-lignin blends in specific ionic liquids. Carbohydrate Polymer Technologies and Applications. 2021;2:100041.

14. Abu-Omar MM, Ford PC. The lignin challenge in catalytic conversion of biomass solids to chemicals and fuels. RSC Sustainability. 2023;1(7):1686–703.

15. Ralph J, Lapierre C, Boerjan W. Lignin structure and its engineering. Current Opinion in Biotechnology. 2019;56:240–9.

16. Eswaran Scd, Subramaniam S, Sanyal U, Rallo R, Zhang X. Molecular structural dataset of lignin macromolecule elucidating experimental structural compositions. Scientific Data. 2022;9(1):647.

17. Vanholme R, Demedts B, Morreel K, Ralph J, Boerjan W. Lignin Biosynthesis and Structure. Plant Physiology. 2010;153(3):895–905.

18. Boerjan W, Ralph J, Baucher M. Lignin Biosynthesis. Annual Review of Plant Biology. 2003;54(Volume 54, 2003):519–46.

19. Vermaas JV, Petridis L, Qi X, Schulz R, Lindner B, Smith JC. Mechanism of lignin inhibition of enzymatic biomass deconstruction. Biotechnology for Biofuels. 2015;8(1):217.

20. Hackenstrass K, Hasani M, Wohlert M. Structure, flexibility and hydration properties of lignin dimers studied with Molecular Dynamics simulations. 2024;78(2):98–108.

21. Vermaas JV, Dixon RA, Chen F, Mansfield SD, Boerjan W, Ralph J, et al. Passive membrane transport of lignin-related compounds. Proceedings of the National Academy of Sciences. 2019;116(46):23117–23.

22. Petridis L, Smith JC. A molecular mechanics force field for lignin. Journal of Computational Chemistry. 2009;30(3):457–67.

23. Vermaas JV, Petridis L, Ralph J, Crowley MF, Beckham GT. Systematic parameterization of lignin for the CHARMM force field. Green Chemistry. 2019;21(1):109–22.

24. D.A. Case HMA, K. Belfon, I.Y. Ben-Shalom, J.T. Berryman, S.R. Brozell, D.S. Cerutti, T.E. Cheatham, III, G.A. Cisneros, V.W.D. Cruzeiro, T.A. Darden, N. Forouzesh, G. Giambaşu, T. Giese, M.K. Gilson, H. Gohlke, A.W. Goetz, J. Harris, S. Izadi, S.A. Izmailov, K. Kasavajhala, M.C. Kaymak, E. King, A. Kovalenko, T. Kurtzman, T.S. Lee, P. Li, C. Lin, J. Liu, T. Luchko, R. Luo, M. Machado, V. Man, M. Manathunga, K.M. Merz, Y. Miao, O. Mikhailovskii, G. Monard, H. Nguyen, K.A. O’Hearn, A. Onufriev, F. Pan, S. Pantano, R. Qi, A. Rahnamoun, D.R. Roe, A. Roitberg, C. Sagui, S. Schott-Verdugo, A. Shajan, J. Shen, C.L. Simmerling, N.R. Skrynnikov, J. Smith, J. Swails, R.C. Walker, J. Wang, J. Wang, H. Wei, X. Wu, Y. Wu, Y. Xiong, Y. Xue, D.M. York, S. Zhao, Q. Zhu, and P.A. Kollman. Amber 2023. 2023.

25. Lahive CW, Kamer PCJ, Lancefield CS, Deuss PJ. An Introduction to Model Compounds of Lignin Linking Motifs; Synthesis and Selection Considerations for Reactivity Studies. ChemSusChem. 2020;13(17):4238–65.

26. Wang J, Wolf RM, Caldwell JW, Kollman PA, Case DA. Development and testing of a general amber force field. Journal of Computational Chemistry. 2004;25(9):1157–74.

27. Grimme S, Bohle F, Hansen A, Pracht P, Spicher S, Stahn M. Efficient Quantum Chemical Calculation of Structure Ensembles and Free Energies for Nonrigid Molecules. The Journal of Physical Chemistry A. 2021;125(19):4039–54.

28. Pracht P, Bohle F, Grimme S. Automated exploration of the low-energy chemical space with fast quantum chemical methods. Physical Chemistry Chemical Physics. 2020;22(14):7169–92.

29. Tian C, Kasavajhala K, Belfon KAA, Raguette L, Huang H, Migues AN, et al. ff19SB: Amino-Acid-Specific Protein Backbone Parameters Trained against Quantum Mechanics Energy Surfaces in Solution. Journal of Chemical Theory and Computation. 2020;16(1):528–52.

30. Kirschner KN, Yongye AB, Tschampel SM, González-Outeiriño J, Daniels CR, Foley BL, et al. GLYCAM06: A generalizable biomolecular force field. Carbohydrates. Journal of Computational Chemistry. 2008;29(4):622–55.

31. Dickson CJ, Walker RC, Gould IR. Lipid21: Complex Lipid Membrane Simulations with AMBER. Journal of Chemical Theory and Computation. 2022;18(3):1726–36.

32. Ivani I, Dans PD, Noy A, Pérez A, Faustino I, Hospital A, et al. Parmbsc1: a refined force field for DNA simulations. Nature Methods. 2016;13(1):55–8.

33. Zgarbová M, Šponer J, Jurečka P. Z-DNA as a Touchstone for Additive Empirical Force Fields and a Refinement of the Alpha/Gamma DNA Torsions for AMBER. Journal of Chemical Theory and Computation. 2021;17(10):6292–301.

34. Zhao Y, Shakeel U, Saif Ur Rehman M, Li H, Xu X, Xu J. Lignin-carbohydrate complexes (LCCs) and its role in biorefinery. Journal of Cleaner Production. 2020;253:120076.

35. Huang C, Jiang X, Shen X, Hu J, Tang W, Wu X, et al. Lignin-enzyme interaction: A roadblock for efficient enzymatic hydrolysis of lignocellulosics. Renewable and Sustainable Energy Reviews. 2022;154:111822.

36. Melro E, Alves L, Antunes FE, Medronho B. A brief overview on lignin dissolution. Journal of Molecular Liquids. 2018;265:578–84.

37. Dixon RA, Barros J. Lignin biosynthesis: old roads revisited and new roads explored. Open Biology. 2019;9(12).

38. Lan W, Lu F, Regner M, Zhu Y, Rencoret J, Ralph SA, et al. Tricin, a Flavonoid Monomer in Monocot Lignification Plant Physiology. 2015;167(4):1284–95.

39. Zhang L, Gellerstedt G. NMR observation of a new lignin structure, a spiro-dienone. Chemical Communications. 2001(24):2744–5.

40. Carlos del Río J, Rencoret J, Gutiérrez A, Kim H, Ralph J. Hydroxystilbenes Are Monomers in Palm Fruit Endocarp Lignins Plant Physiology. 2017;174(4):2072–82.

41. Ralph J, Lapierre C, Lu F, Marita JM, Pilate G, Van Doorsselaere J, et al. NMR Evidence for Benzodioxane Structures Resulting from Incorporation of 5-Hydroxyconiferyl Alcohol into Lignins of O-Methyltransferase-Deficient Poplars. Journal of Agricultural and Food Chemistry. 2001;49(1):86–91.

42. Karhunen P, Rummakko P, Sipilä J, Brunow G, Kilpeläinen I. The formation of dibenzodioxocin structures by oxidative coupling. A model reaction for lignin biosynthesis. Tetrahedron Letters. 1995;36(25):4501–4.

43. Yuan T-Q, Sun S-N, Xu F, Sun R-C. Characterization of Lignin Structures and Lignin– Carbohydrate Complex (LCC) Linkages by Quantitative 13C and 2D HSQC NMR Spectroscopy. Journal of Agricultural and Food Chemistry. 2011;59(19):10604–14.

44. Lu F, Karlen SD, Regner M, Kim H, Ralph SA, Sun R-C, et al. Naturally p-Hydroxybenzoylated Lignins in Palms. BioEnergy Research. 2015;8(3):934–52.

45. Ralph J, Hatfield RD, Quideau S, Helm RF, Grabber JH, Jung H-JG. Pathway of p-Coumaric Acid Incorporation into Maize Lignin As Revealed by NMR. Journal of the American Chemical Society. 1994;116(21):9448–56.

46. Giummarella N, Pu Y, Ragauskas AJ, Lawoko M. A critical review on the analysis of lignin carbohydrate bonds. Green Chemistry. 2019;21(7):1573–95.

47. Landrum G. RDKit: Open-source cheminformatics. https://www.rdkit.org (accessed August 28th, 2024). 2010.

48. Bannwarth C, Ehlert S, Grimme S. GFN2-xTB—An Accurate and Broadly Parametrized Self-Consistent Tight-Binding Quantum Chemical Method with Multipole Electrostatics and Density-Dependent Dispersion Contributions. Journal of Chemical Theory and Computation. 2019;15(3):1652–71.

49. Bannwarth C, Caldeweyher E, Ehlert S, Hansen A, Pracht P, Seibert J, et al. Extended tight-binding quantum chemistry methods. WIREs Computational Molecular Science. 2021;11(2):e1493.

50. Ehlert S, Stahn M, Spicher S, Grimme S. Robust and Efficient Implicit Solvation Model for Fast Semiempirical Methods. Journal of Chemical Theory and Computation. 2021;17(7):4250–61.

51. Spicher S, Grimme S. Robust Atomistic Modeling of Materials, Organometallic, and Biochemical Systems. Angewandte Chemie International Edition. 2020;59(36):15665–73.

52. Bayly CI, Cieplak P, Cornell W, Kollman PA. A well-behaved electrostatic potential based method using charge restraints for deriving atomic charges: the RESP model. The Journal of Physical Chemistry. 1993;97(40):10269–80.

53. Vanquelef E, Simon S, Marquant G, Garcia E, Klimerak G, Delepine JC, et al. R.E.D. Server: a web service for deriving RESP and ESP charges and building force field libraries for new molecules and molecular fragments. Nucleic Acids Research. 2011;39(suppl_2):W511–W7.

54. Wang FB, J.-P.; Cieplak, P.; Dupradeau, F.-Y. R.E.D. Server Development - Performing calculations with the PyRED program: Application to charge derivation, force field library building and force field parameter generation 2014 [Available from: https://upjv.q4md-forcefieldtools.org/Tutorial/Tutorial-4.php.

55. Ditchfield R, Hehre WJ, Pople JA. Self-Consistent Molecular-Orbital Methods. IX. An Extended Gaussian-Type Basis for Molecular-Orbital Studies of Organic Molecules. The Journal of Chemical Physics. 1971;54(2):724–8.

56. Hehre WJ, Ditchfield R, Pople JA. Self—Consistent Molecular Orbital Methods. XII. Further Extensions of Gaussian—Type Basis Sets for Use in Molecular Orbital Studies of Organic Molecules. The Journal of Chemical Physics. 1972;56(5):2257–61.

57. Slater JC. A Simplification of the Hartree-Fock Method. Physical Review. 1951;81(3):385–90.

58. Dunning TH, Jr. Gaussian basis sets for use in correlated molecular calculations. I. The atoms boron through neon and hydrogen. The Journal of Chemical Physics. 1989;90(2):1007–23.

59. Møller C, Plesset MS. Note on an Approximation Treatment for Many-Electron Systems. Physical Review. 1934;46(7):618–22.

60. Becke AD. Density-functional thermochemistry. III. The role of exact exchange. The Journal of Chemical Physics. 1993;98(7):5648–52.

61. Chai J-D, Head-Gordon M. Long-range corrected hybrid density functionals with damped atom–atom dispersion corrections. Physical Chemistry Chemical Physics. 2008;10(44):6615–20.

62. Frisch MJ, Trucks GW, Schlegel HB, Scuseria GE, Robb MA, Cheeseman JR, et al. Gaussian 16 Rev. C.01. 2016.

63. Dupradeau F-Y, Pigache A, Zaffran T, Savineau C, Lelong R, Grivel N, et al. The R.E.D. tools: advances in RESP and ESP charge derivation and force field library building. Physical Chemistry Chemical Physics. 2010;12(28):7821–39.

64. Zhao S, Wei H, Cieplak P, Duan Y, Luo R. PyRESP: A Program for Electrostatic Parameterizations of Additive and Induced Dipole Polarizable Force Fields. Journal of Chemical Theory and Computation. 2022;18(6):3654–70.

65. Lapsien M, Bonus M, Gahan L, Raguin A, Gohlke H. PyPE_RESP: A Tool to Facilitate and Standardize Derivation of RESP Charges. Journal of Chemical Information and Modeling. 2025;65(9):4251–6.

66. Grimme S, Hansen A, Ehlert S, Mewes J-M. r2SCAN-3c: A “Swiss army knife” composite electronic-structure method. The Journal of Chemical Physics. 2021;154(6).

67. Neese F. Software update: The ORCA program system—Version 5.0. WIREs Computational Molecular Science. 2022;12(5):e1606.

68. Caldeweyher E, Ehlert S, Hansen A, Neugebauer H, Spicher S, Bannwarth C, et al. A generally applicable atomic-charge dependent London dispersion correction. The Journal of Chemical Physics. 2019;150(15).

69. Chai J-D, Head-Gordon M. Systematic optimization of long-range corrected hybrid density functionals. The Journal of Chemical Physics. 2008;128(8).

70. Weigend F, Ahlrichs R. Balanced basis sets of split valence, triple zeta valence and quadruple zeta valence quality for H to Rn: Design and assessment of accuracy. Physical Chemistry Chemical Physics. 2005;7(18):3297–305.

71. Lange RT. evosax: JAX-Based Evolution Strategies. Proceedings of the Companion Conference on Genetic and Evolutionary Computation; Lisbon, Portugal: Association for Computing Machinery; 2023. p. 659–62.

72. Bradbury J, Frostig R, Hawkins P, Johnson MJ, Leary C, Maclaurin D, et al. JAX: composable transformations of Python+NumPy programs. 0.3.13 ed2018.

73. Le Grand S, Götz AW, Walker RC. SPFP: Speed without compromise—A mixed precision model for GPU accelerated molecular dynamics simulations. Computer Physics Communications. 2013;184(2):374–80.

74. Roe DR, Cheatham TE, III. PTRAJ and CPPTRAJ: Software for Processing and Analysis of Molecular Dynamics Trajectory Data. Journal of Chemical Theory and Computation. 2013;9(7):3084–95.

75. Groom CR, Bruno IJ, Lightfoot MP, Ward SC. The Cambridge Structural Database. Acta Crystallographica Section B. 2016;72(2):171–9.

76. Loncharich RJ, Brooks BR, Pastor RW. Langevin dynamics of peptides: The frictional dependence of isomerization rates of N-acetylalanyl-N′-methylamide. Biopolymers. 1992;32(5):523–35.

77. Berendsen HJC, Postma JPM, Van Gunsteren WF, Dinola A, Haak JR. Molecular dynamics with coupling to an external bath. The Journal of Chemical Physics. 1984;81(8):3684–90.

78. Martínez L, Andrade R, Birgin EG, Martínez JM. PACKMOL: A package for building initial configurations for molecular dynamics simulations. Journal of Computational Chemistry. 2009;30(13):2157–64.

79. Kräutler V, Van Gunsteren WF, Hünenberger PH. A fast SHAKE algorithm to solve distance constraint equations for small molecules in molecular dynamics simulations. Journal of Computational Chemistry. 2001;22(5):501–8.

80. Kaus JW, Pierce LT, Walker RC, McCammon JA. Improving the Efficiency of Free Energy Calculations in the Amber Molecular Dynamics Package. Journal of Chemical Theory and Computation. 2013;9(9):4131–9.

81. Jorgensen WL, Chandrasekhar J, Madura JD, Impey RW, Klein ML. Comparison of simple potential functions for simulating liquid water. The Journal of Chemical Physics. 1983;79(2):926–35.

82. Wu Z, Dotson DL, Alibay I, Allen BK, Barhaghi MS, Hénin J, et al. alchemlyb: the simple alchemistry library. Journal of Open Source Software. 2024;9(101):6934.

83. Mobley DL, Guthrie JP. FreeSolv: a database of experimental and calculated hydration free energies, with input files. Journal of Computer-Aided Molecular Design. 2014;28(7):711–20.

84. Grimme S. Semiempirical GGA-type density functional constructed with a long-range dispersion correction. Journal of Computational Chemistry. 2006;27(15):1787–99.

85. Caleman C, van Maaren PJ, Hong M, Hub JS, Costa LT, van der Spoel D. Force Field Benchmark of Organic Liquids: Density, Enthalpy of Vaporization, Heat Capacities, Surface Tension, Isothermal Compressibility, Volumetric Expansion Coefficient, and Dielectric Constant. Journal of Chemical Theory and Computation. 2012;8(1):61–74.

86. Kashefolgheta S, Wang S, Acree WE, Hünenberger PH. Evaluation of nine condensed-phase force fields of the GROMOS, CHARMM, OPLS, AMBER, and OpenFF families against experimental cross-solvation free energies. Physical Chemistry Chemical Physics. 2021;23(23):13055–74.

87. Wang J, Hou T. Application of Molecular Dynamics Simulations in Molecular Property Prediction. 1. Density and Heat of Vaporization. Journal of Chemical Theory and Computation. 2011;7(7):2151–65.

88. Izadi S, Anandakrishnan R, Onufriev AV. Building Water Models: A Different Approach. The Journal of Physical Chemistry Letters. 2014;5(21):3863–71.

