## Supporting Information for "LignAmb25: A Comprehensive AMBER Force Field Addressing Lignin’s Structural and Chemical Diversity"

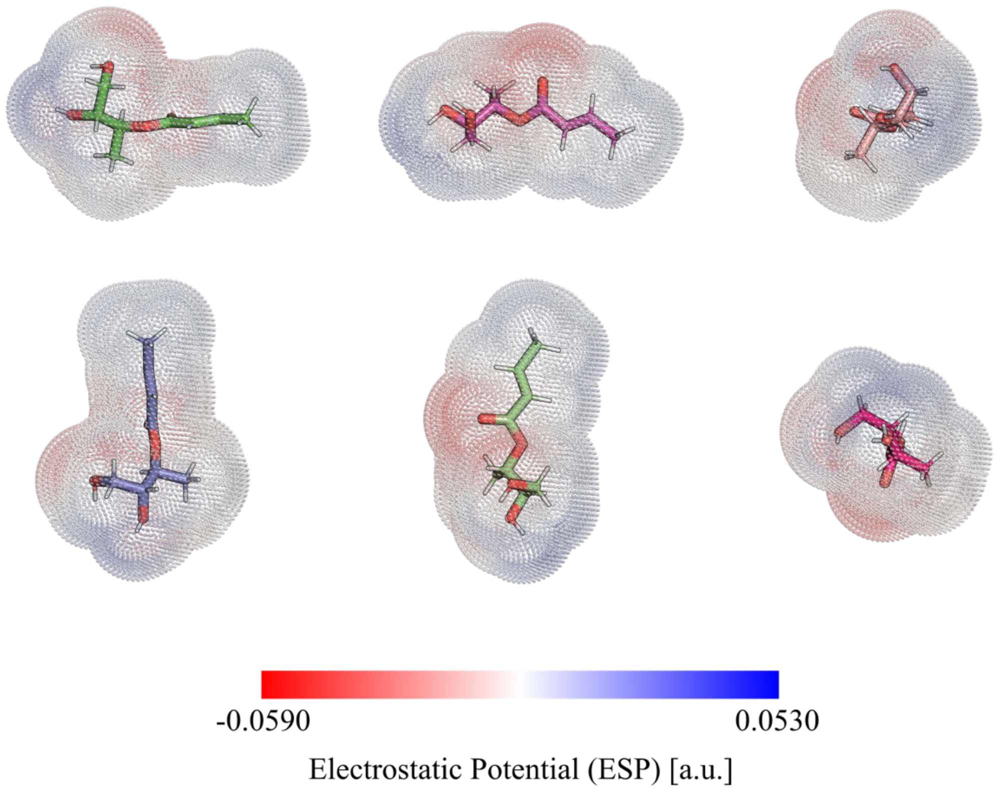


**Figure S1: Example for orientation sampling performed for RESP Fitting.** Orientations from top left to bottom right: Initial orientation, 90° rotation around x-axis, 90° rotation around y-axis, 90° rotation around z-axis, 90° rotation around x-axis + 90° rotation around z-axis, 90° rotation around y-axis + 90° rotation around z-axis.

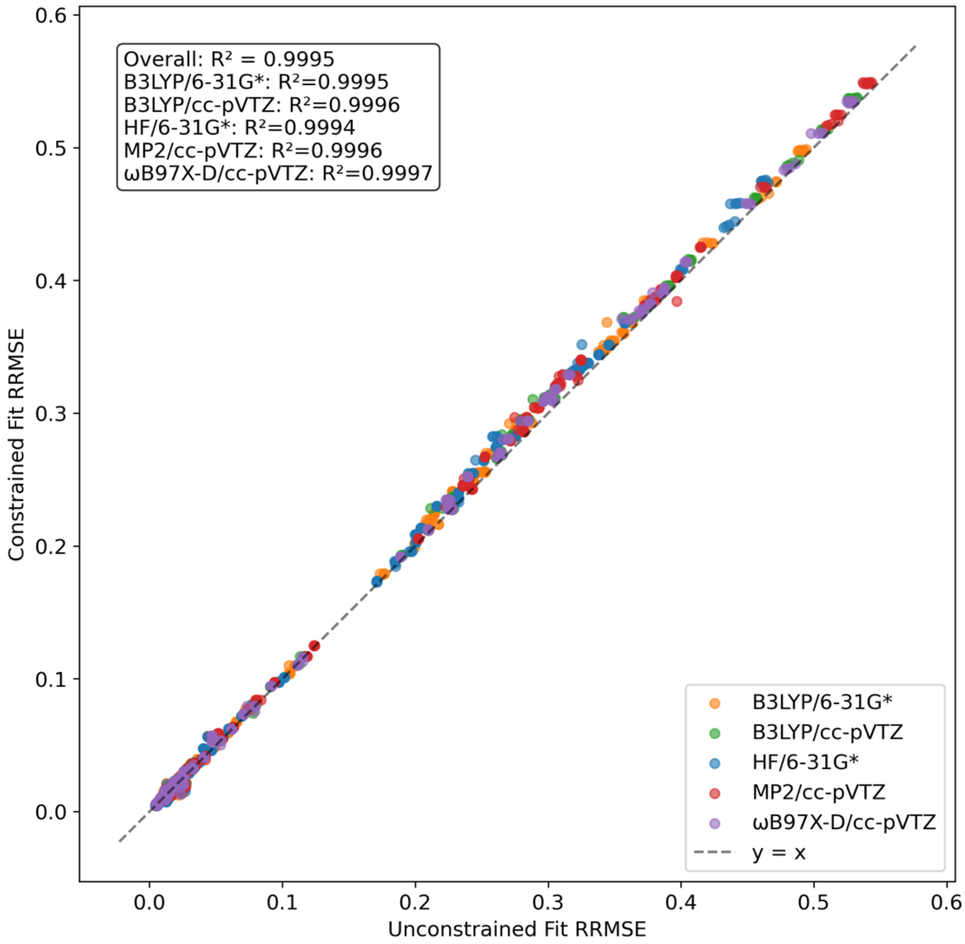


**Figure S2: Correlation between RRMSE of RESP Fits with and without cap constraints.**

Combined results over all RESP charge restraint weights per level of theory studied are shown.

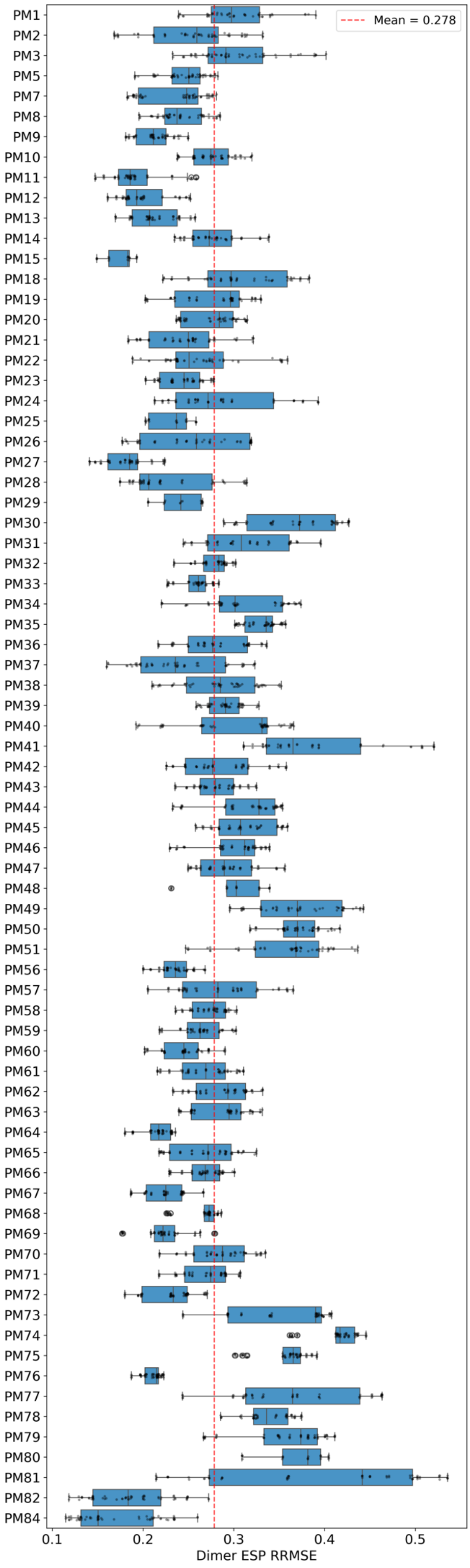
Mean = 0.278
PM1
PM2
PM3
PM5
PM7
PM8
PM9
PM10
PM11
PM12
PM13
PM14
PM15
PM18
PM19
PM20
PM21
PM22
PM23
PM24
PM25
PM26
PM27
PM28
PM29
PM30
PM31
PM32
PM33
PM34
PM35
PM36
PM37
PM38
PM39
PM40
PM41
PM42
PM43
PM44
PM45
PM46
PM47
PM48
PM49
PM50
PM51
PM56
PM57
PM58
PM59
PM60
PM61
PM62
PM63
PM64
PM65
PM66
PM67
PM68
PM69
PM70
PM71
PM72
PM73
PM74
PM75
PM76
PM77
PM78
PM79
PM80
PM81
PM82
PM84
0.1
0.2
0.3
0.4
0.5
Dimer ESP RRMSE

**Figure S3: RRMSE distribution of ESP reproduction of lignin multimers based on force field fragment RESP charges.** Data was taken for each analyzed charge restraint weight of all levels of theory studied.

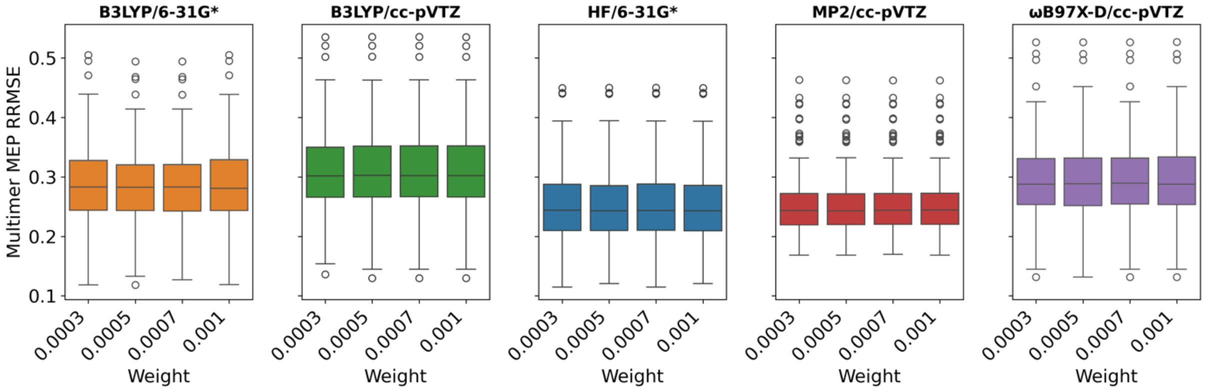


**Figure S4: Comparison of ESP reproduction for dimers from RESP charges for different charge restraint weights and levels of theory.**

### Interunit linkages

A

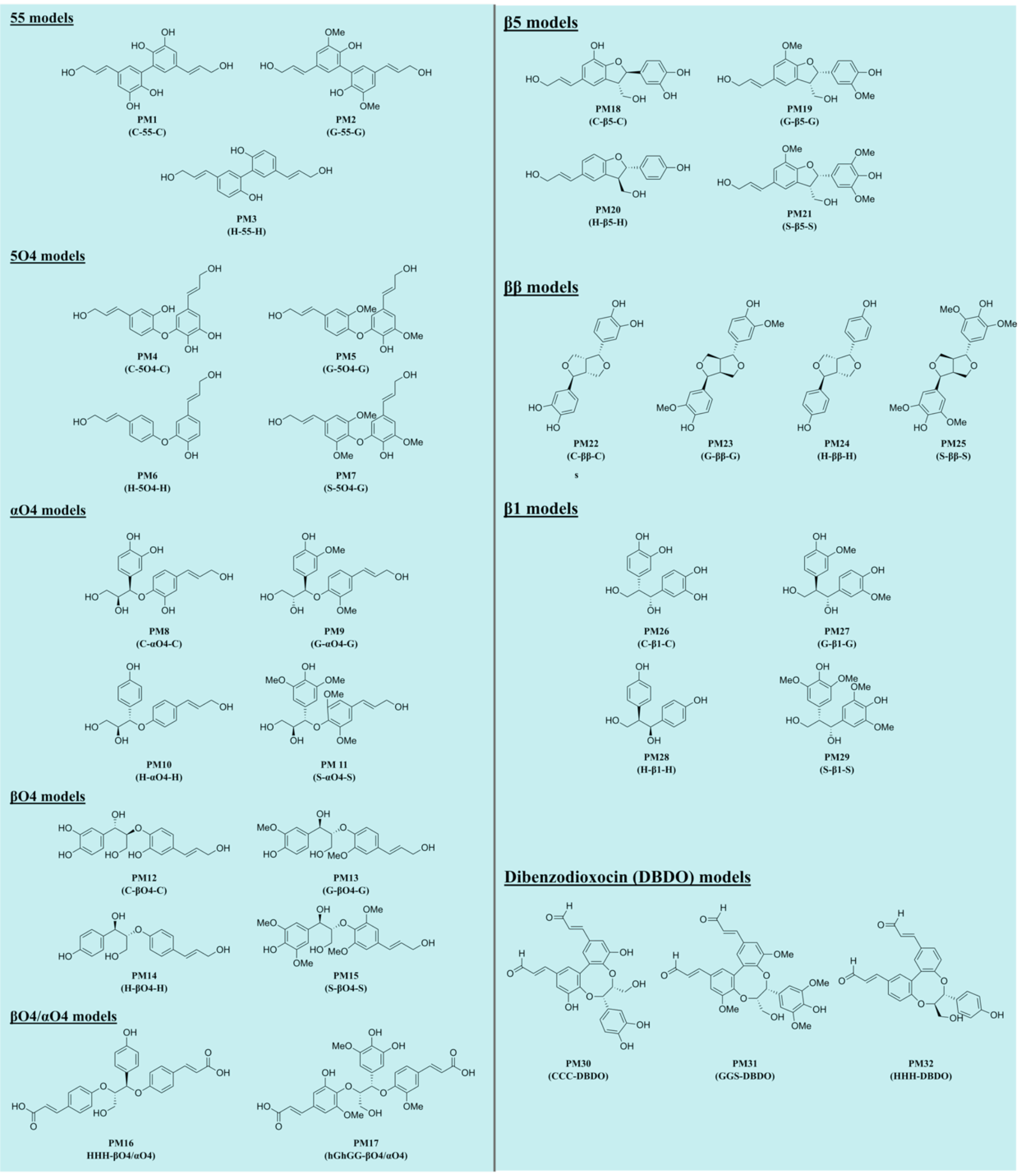

### Hydroxycinnamic acid conjugates

**B**

**α-Cinnamate (αCM) models**

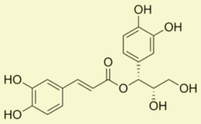

**PM33**
**(C-αCM-C)**

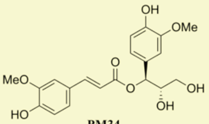

**PM34**
**(G-αCM-G)**

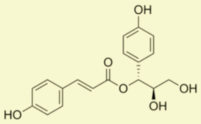

**PM35**
**(H-αCM-H)**

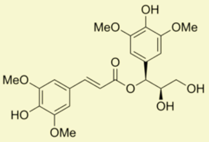

**PM36**
**(S-αCM-S)**

**γ-Cinnamate (γCM) models**

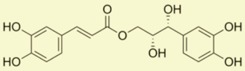

**PM37**
**(C-γCM-C)**

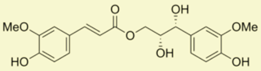

**PM38**
**(G-γCM-G)**

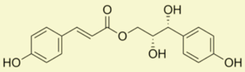

**PM39**
**(H-γCM-H)**

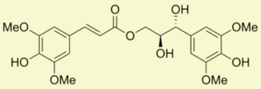

**PM40**
**(S-γCM-S)**

**α-Benzoate (αBZ) models**

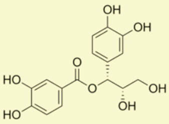

**PM41**
**(C-αBZ-C)**

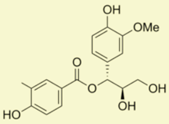

**PM42**
**(G-αBZ-G)**

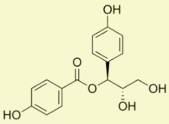

**PM43**
**(H-αBZ-H)**

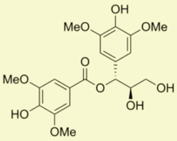

**PM44**
**(S-αBZ-S)**

**γ-Benzoate (γBZ) models**

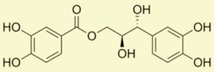

**PM45**
**(C-γBZ-C)**

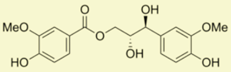

**PM46**
**(G-γBZ-G)**

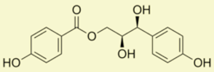

**PM47**
**(H-γBZ-H)**

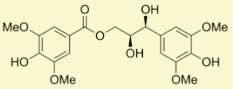

**PM48**
**(S-γBZ-S)**

**Diferulate (DF) models**

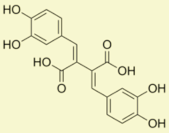

**PM49**
**(C-DF-C)**

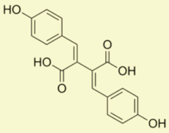

**PM50**
**(H-DF-H)**

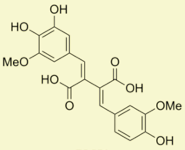

**PM51**
**(hG-DF-G)**

**Cyclic diferulate (cycDF) models**

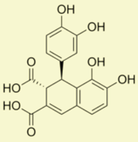

**PM52**
**(C-cycDF-C)**

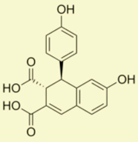

**PM53**
**(H-cycDF-H)**

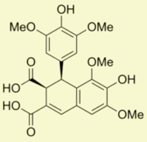

**PM54**
**(S-cycDF-S)**

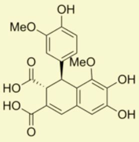

**PM55**
**(hG-cycDF-G)**

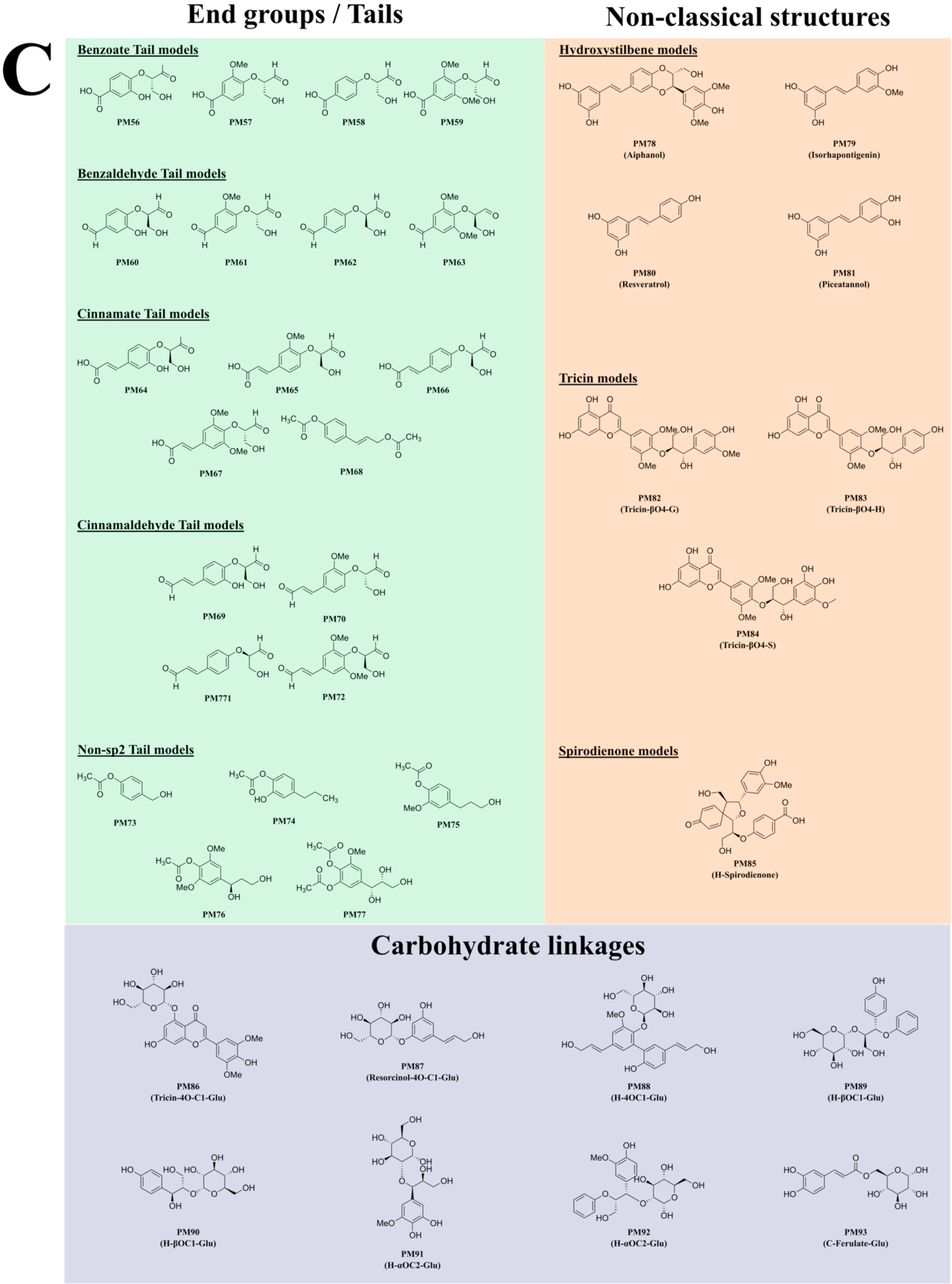
C
End groups / Tails
Non-classical structures
Benzoate Tail models
PM56
PM57
PM58
PM59
Benzaldehyde Tail models
PM60
PM61
PM62
PM63
Cinnamate Tail models
PM64
PM65
PM66
PM67
PM68
Cinnamaldehyde Tail models
PM69
PM70
PM771
PM72
Non-sp2 Tail models
PM73
PM74
PM75
PM76
PM77
Hydroxystilbene models
PM78 (Aiphanol)
PM79 (Isorhapontigenin)
PM80 (Resveratrol)
PM81 (Piceatannol)
Tricin models
PM82 (Tricin-βO4-G)
PM83 (Tricin-βO4-H)
PM84 (Tricin-βO4-S)
Spirodienone models
PM85 (H-Spirodienone)
Carbohydrate linkages
PM86 (Tricin-4O-C1-Glu)
PM87 (Resorcinol-4O-C1-Glu)
PM88 (H-4OC1-Glu)
PM89 (H-βOC1-Glu)
PM90 (H-βOC1-Glu)
PM91 (H-αOC2-Glu)
PM92 (H-αOC2-Glu)
PM93 (C-Ferulate-Glu)

**Figure S5: Parameterization models covering end groups, carbohydrate linkages, and less commonly encountered lignin monomers.** A: Models covering commonly encountered lignin linkages. B: Models covering hydroxycinnamic acid conjugates encountered in lignin polymers. C: Models covering end groups, carbohydrate linkages, and less commonly encountered lignin monomers. All models except for lignin-carbohydrate models were used for charge transferability assessment due to GLYCAM charges being predetermined in the latter case.

**Table S1: Overview of atom types included in the LignAmb25 force field**

| Atom type | Target atoms |
|---|---|
| Ca | Carbonyl carbon |
| Cb | Beta carbon in propyl side chain |
| Cd | Alpha carbon bonded to aromatic ring |
| CD | Alpha carbon bonded to aromatic ring (alternate) |
| Ce | Alpha carbon with ether linkage |
| Cf | Gamma carbon in β-O-4 linkage (from torsion fit) |
| Ci | Alpha carbon in β-5 linkage (from torsion fit) |
| Cm | Methoxy carbon |
| Co | Alpha carbon with hydroxyl group |
| Cr | Gamma carbon in side chain |
| Ct | Gamma carbon in side chain (alternate) |
| Cx | Carbonyl carbon (alternate) |
| cE | Aromatic carbon with ether substituent |
| cH | Aromatic carbon with H (unsubstituted) |
| cM | Aromatic carbon with methoxy group |
| cm | Aromatic carbon with methoxy group (alternate) |
| cN | Aromatic carbon at C1 |
| cO | Aromatic carbon with hydroxyl group |
| HR | Aromatic ring hydrogen |
| Hd | Hydrogen on alpha carbon |
| Hm | Methoxy hydrogen |
| Hq | Hydroxyl hydrogen |

| | |
|---|---|
| Hr | Hydrogen on beta/gamma carbon |
| Hx | Hydrogen on carbon adjacent to carbonyl |
| OE | Ether oxygen in aryl ether linkage |
| Ob | Bridging ether oxygen (β-O-4 linkage) |
| Oc | Phenolic/aliphatic hydroxyl oxygen |
| Oe | Ether oxygen (aliphatic C-O-C) |
| Om | Methoxy oxygen |
| Or | Ring ether oxygen (in furan/dioxane) |
| Oz | Carbonyl oxygen |

**Table S2: Details on CENSO settings used for identification of representative conformers from CREST-generated ensembles.** All CENSO parts were run using ORCA 6.0 as the QM backend.

| Stage | Functional/Basis | Solvation model | Energy Threshold |
|---|---|---|---|
| Prescreening (Part 0) | pbe-d4/def2-SV(P) | – | 5.0 kcal·mol$^{-1}$ |
| Screening (Part 1) | r2scan-3c/def2-TZVP | SMD | 3.5 kcal·mol$^{-1}$ |
| Optimization (Part 2) | r2scan-3c/def2-TZVP | SMD | 3.0 kcal·mol$^{-1}$ |
| Refinement (Part 3) | wb97x-d4/def2-TZVPP | SMD | 95%* |

*Threshold for additive Boltzmann population beyond which conformers will be neglected

**Table S3: Mean RRMSE values obtained for RESP fits at different levels of theory and charge restraint weights.** The RRMSE was calculated over all force field fragments ($n$ = 134).

| Level of theory | Weight | Mean RRMSE | Standard deviation |
|---|---|---|---|
| B3LYP/6-31G* | 0.0003 | 0.08527 | 0.13230 |
| | 0.0005 | 0.08530 | 0.13228 |
| | 0.0007 | 0.08525 | 0.13234 |
| | 0.001 | 0.08520 | 0.13247 |
| B3LYP/cc-pVTZ | 0.0003 | 0.09213 | 0.14215 |
| | 0.0005 | 0.09220 | 0.14214 |
| | 0.0007 | 0.09224 | 0.14218 |
| | 0.001 | 0.09221 | 0.14228 |
| HF/6-31G* | 0.0003 | 0.08104 | 0.12664 |
| | 0.0005 | 0.08086 | 0.12675 |
| | 0.0007 | 0.08101 | 0.12666 |
| | 0.001 | 0.08094 | 0.12682 |
| MP2/cc-pVTZ | 0.0003 | 0.09522 | 0.14683 |
| | 0.0005 | 0.09450 | 0.14659 |
| | 0.0007 | 0.09528 | 0.14683 |
| | 0.001 | 0.09531 | 0.14682 |
| wB97X-D/cc-pVTZ | 0.0003 | 0.09143 | 0.14159 |
| | 0.0005 | 0.09146 | 0.14158 |
| | 0.0007 | 0.09145 | 0.14160 |
| | 0.001 | 0.09142 | 0.14168 |

**Table S4: Details on ORCA settings used for potential energy surface scans of bond stretching and bond angle bending**

| | | |
|---|---|---|
| **SCF Convergence (tightscf)** | TolE | 1e-8 |
| | TolMAXP | 1e-7 |
| | TolRMSP | 5e-9 |
| | TolErr | 5e-7 |
| | Thresh | 2.5e-11 |
| | TCut | 2.5e-12 |
| | DFTGrid.BFCut | 1e-11 |
| | TolG | 1e-5 |
| | TolX | 1e-5 |
| | Z_Tol | 1e-4 |
| | Maxiter | 1000 |
| **Optimization convergence (tightopt)** | TolE | 1e-6 |
| | TolRMSG | 3e-5 |
| | TolMaxG | 1e-4 |
| | TolRMSD | 6e-4 |
| | TolMaxD | 1e-3 |
| | Maxiter | 1000 |

**Table S5: Details on ORCA settings used for potential energy surface scans of torsion angles.** The optimization convergence criteria were less strict to reduce computational costs.

| | | |
|---|---|---|
| **SCF Convergence (tightscf)** | TolE | 1e-8 |
| | TolMAXP | 1e-7 |
| | TolRMSP | 5e-9 |
| | TolErr | 5e-7 |
| | Thresh | 2.5e-11 |
| | TCut | 2.5e-12 |
| | DFTGrid.BFCut | 1e-11 |
| | TolG | 1e-5 |
| | TolX | 1e-5 |
| | Z_Tol | 1e-4 |
| | Maxiter | 1000 |
| **Optimization convergence (normalopt)** | TolE | 5e-6 |
| | TolRMSG | 1e-4 |
| | TolMaxG | 3e-4 |
| | TolRMSD | 2e-3 |
| | TolMaxD | 4e-3 |
| | Maxiter | 1000 |

**Table S6: Comparison of mean crystal properties after simulation using the established lignin forcefield for CHARMM (CHARMM), the GAFF2 force field parameters with LignAmb25 RESP charges (GAFF2), and the MP2/cc-pVTZ-based LignAmb25 force field (LignAmb25).**

| Name (CSD Code) | ρ ratio $\left(\frac{\rho_{sim}}{\rho_{exp}}\right)$[a] | | | RMSD$^C$ [Å][b] | | | RMSD$^M$ [Å][c] | | |
|---|---|---|---|---|---|---|---|---|---|
| | LignAmb[d] | CHARMM | GAFF2 | LignAmb | CHARMM | GAFF2 | LignAmb | CHARMM | GAFF2 |
| Catechol (CATCOL13) | 1.0657(2) | 0.9687(2) | 1.0603(3) | 0.370(7) | 0.977(8) | 0.411(8) | 0.04(0) | 0.05(1) | 0.04(0) |
| *p*-Hydroxy benzaldehyde (PHBALD11) | 1.0240(8) | 0.9755(6) | 1.0504(8) | 1.056(1) | 1.09(2) | 0.937(1) | 0.10(0) | 0.11(3) | 0.10(0) |
| Vanilin (YUHTEA01) | 1.0398(3) | 0.9749(2) | 1.0662(4) | 0.732(8) | 1.331(8) | 0.542(1) | 0.09(0) | 0.10(2) | 0.09(0) |
| Vanilin (YUHTEA03) | 1.0312(7) | 1.061(1) | 1.0591(7) | 0.911(2) | 1.52(3) | 0.716(2) | 0.12(0) | 0.14(4) | 0.13(0) |
| Siringaldehyde (IZALAW) | 0.9183(7) | 0.9491(4) | 0.9225(7) | 0.904(1) | 8.92(1) | 0.897(1) | 0.14(0) | 0.19(4) | 0.19(0) |
| Coniferylaldehyde (SIPKEH) | 0.9544(7) | 0.9913(6) | 0.9698(9) | 0.632(1) | 3.16(2) | 0.635(2) | 0.17(0) | 0.5(1) | 0.19(0) |
| Vanilic acid (CEHGUS) | 0.9621(8) | 0.9360(4) | 0.9366(8) | 0.682(1) | 1.01(1) | 0.808(1) | 0.15(0) | 0.15(4) | 0.15(0) |
| Ferulic acid (GASVOL01) | 0.9824(3) | 0.9557(2) | 1.0042(3) | 0.322(4) | 0.837(6) | 0.276(3) | 0.14(0) | 0.11(3) | 0.14(0) |
| G-βO4-G (RABWUM) | 0.9950(3) | 0.9630(2) | 0.9948(4) | 1.367(2) | 0.592(7) | 0.310(5) | 0.37(1) | 0.20(4) | 0.15(0) |
| G-βO4-G (SIPPEM) | 0.9613(6) | 0.9538(3) | 0.9704(5) | 0.943(4) | 0.93(1) | 0.526(9) | 0.33(1) | 0.36(9) | 0.23(0) |
| S-βO4-G (VADDOT) | 1.0386(7) | 0.9546(3) | 1.0498(6) | 0.671(1) | 1.15(1) | 0.579(1) | 0.30(0) | 0.29(7) | 0.26(0) |
| S-βO4-S (SAZHEG) | 0.9899(1) | 0.9474(4) | 1.0062(9) | 0.988(1) | 0.95(1) | 0.561(2) | 0.53(0) | 0.25(6) | 0.28(1) |
| S-βO4-S (FOCGUA) | 1.0581(2) | 0.9559(1) | 1.0628(3) | 0.488(7) | 0.917(7) | 0.463(7) | 0.17(0) | 0.19(4) | 0.21(0) |
| S-βO4-S (IDIKIP) | 1.0138(6) | 0.9480(2) | 1.0301(4) | 1.185(8) | 0.84(1) | 0.568(9) | 0.39(1) | 0.19(4) | 0.25(0) |
| G-ββ-G (INELIW) | 0.9814(4) | 0.9626(2) | 0.9591(1) | 0.433(7) | 0.786(9) | 1.686(5) | 0.14(0) | 0.11(7) | 0.53(1) |
| G-ββ-G | 0.9882(4) | 0.9656(2) | 0.9771(4) | 0.904(4) | 0.82(2) | 0.971(6) | 0.11(0) | 0.12(9) | 0.44(0) |

| | | | | | | | | | |
|---|---|---|---|---|---|---|---|---|---|
| (INELIW01) | | | | | | | | | |
| G-ββ-G (FAFXUF) | 0.9744(7) | 0.9368(4) | 0.8976(2) | 0.440(9) | 0.99(1) | 3.365(8) | 0.21(0) | 0.19(4) | 1.45(2) |
| G-β5-G (FUMVUE) | 1.0117(7) | 0.9357(3) | 1.0226(6) | 0.684(1) | 1.01(1) | 0.553(1) | 0.23(0) | 0.25(7) | 0.20(0) |
| DBDO (TUGWAT) | 1.0111(5) | 0.9518(4) | 1.0048(4) | 0.528(5) | 0.88(2) | 0.362(6) | 0.39(0) | 0.29(4) | 0.22(0) |

[a]Density ratio calculated from the simulated ($\rho_{sim}$) and experimentally measured crystal structure ($\rho_{exp}$)

[b]RMSD$^C$ calculated as the RMSD of the simulated relative to the deposited crystal structure

[c]RMSD$^M$ calculated as the average intramolecular RMSD of the molecule in the simulated relative to the deposited crystal structure

[d]Data for GAFF2 and LignAmb25 were obtained from 20 independent replica simulations by taking the last 200 frames from the simulation trajectory, corresponding to the last 10 ns of the total 20 ns simulated, whereas data for CHARMM were taken as reported from the source publication. Uncertainty in the last digit is reported in parentheses.
